# Uncovering causal relationships in single-cell omic studies with causarray

**DOI:** 10.1101/2025.01.30.635593

**Authors:** Jin-Hong Du, Maya Shen, Hansruedi Mathys, Kathryn Roeder

## Abstract

Advances in single-cell sequencing and CRISPR technologies have enabled detailed case-control comparisons and experimental perturbations at single-cell resolution. However, uncovering causal relationships in observational genomic data remains challenging due to selection bias and inadequate adjustment for unmeasured confounders, particularly in heterogeneous datasets. To address these challenges, we introduce causarray, a robust causal inference framework for analyzing array-based genomic data at both pseudo-bulk and single-cell levels under unmeasured confounding. causarray integrates a generalized confounder adjustment method to account for unmeasured confounders and employs semiparametric inference with flexible machine learning techniques to ensure robust statistical estimation of treatment effects. Benchmarking results show that causarray robustly separates treatment effects from confounders while preserving biological signals across diverse settings. We also apply causarray to two single-cell genomic studies: (1) an in vivo Perturb-seq study of autism risk genes in developing mouse brains and (2) a case-control study of Alzheimer’s disease using three human brain transcriptomic datasets. In these applications, causarray identifies clustered causal effects of multiple autism risk genes and consistent causally affected genes across Alzheimer’s disease datasets, uncovering biologically relevant pathways directly linked to neuronal development and synaptic functions that are critical for understanding disease pathology.

## Introduction

The advent of genomic research has revolutionized our understanding of biological systems and disease mechanisms. In particular, advances in single-cell RNA sequencing (scRNA-seq) have provided unprecedented resolution of gene expression at the cellular level, enabling detailed characterization of cellular heterogeneity and its relevance to health and disease (1–4). Likewise, understanding the regulatory circuits that govern the phenotypic landscape of human cells has long been considered a formidable challenge, but recent experimental innovations, such as pooled CRISPR-based perturbation assays, are making this goal increasingly attainable (5).

Fully realizing the potential of these technologies, however, requires analytical frameworks that move beyond mere association to uncover causal relationships at single-cell resolution (6–8). Association studies identify correlations between treatments and outcomes, whereas causal inference seeks to estimate the effect of an intervention on an outcome. A widely used approach for causal inference is the potential outcomes framework, which contrasts observed outcomes with their unobserved counterparts, the counterfactuals, to quantify causal effects (8, 9). Developing such causal models is essential for understanding biological processes and disease mechanisms, with important implications for treatments, precision medicine, genomic medicine, and related fields (10, 11).

One of the primary challenges in leveraging scRNA-seq and CRISPR data for causal inference is the presence of unmeasured confounders due to biological factors, such as correlated gene expression and technical factors such as batch effects. Furthermore, most genomic studies are observational in nature. Unlike randomized controlled trials, observational studies lack complete knowledge of the disease or treatment assignment mechanism, leading to potential biases in counterfactual estimation. In CRISPR screens (5, 12–14), perturbed cells are contrasted with non-targeting controls, but assignment is not fully random: continuous, cell-level variables such as cell size or differential exposure can leak into effect estimates, a problem amplified in vivo where more complex cellular environments exacerbate confounding. Although in vivo CRISPR screens are feasible (15, 16), many screens still rely on a few well-studied cell lines (17), limiting generalizability. As perturbation technologies scale, including multimodal assays (5), the need for robust causal inference that explicitly models unmeasured confounders becomes even more acute.

The presence of unmeasured confounders can undermine the validity of causal conclusions in observational studies (18, 19). Existing methods for causal estimation, such as CoCoA-diff (7) and CINEMA-OT (20), rely on matching techniques that assume the causal structure is transferable between treatment and control groups. However, this assumption breaks down when covariate distributions differ significantly across groups, leading to biased estimates. On the other hand, to adjust for confounding and unwanted variation in statistical inference, other methods like surrogate variable analysis (SVA) (21) and remove unwanted variation (RUV) (22) assume additive relationships between covariates and outcomes via linear models. While effective for certain bulk RNA-seq datasets, these approaches often fail to capture the sparsity, zero inflation, and overdispersion inherent in single-cell genomic data (18, 23). Tackling these challenges requires integrating robust confounder adjustment with flexible modeling techniques to ensure valid causal inference in complex genomic data.

In response to these challenges, we introduce a new framework for applying causal inference in omic studies. Our approach leverages generalized factor models tailored to count data to account for unmeasured confounders, ensuring robust adjustment for unmeasured confounders while preserving biological signals. It further relies on the potential outcomes framework and employs a robust estimation procedure, which combines outcome and propensity score models to ensure reliable statistical inference even if one model is misspecified (24, 25). This framework effectively addresses biases introduced by both observed and unobserved confounders, making it particularly well-suited for analyzing complex genomic data at both pseudo-bulk and single-cell levels (Fig. 1a). By integrating advanced statistical and machine learning techniques with a causal inference framework, our method enables a range of downstream analyses, including accurate estimation of counterfactual distributions, causal gene detection, and conditional treatment effect analysis. This approach not only improves the interpretability and precision of genomic analyses but also uncovers critical insights into gene expression dynamics under disease or perturbation conditions, advancing our understanding of underlying biological mechanisms.

**Fig. 1.**
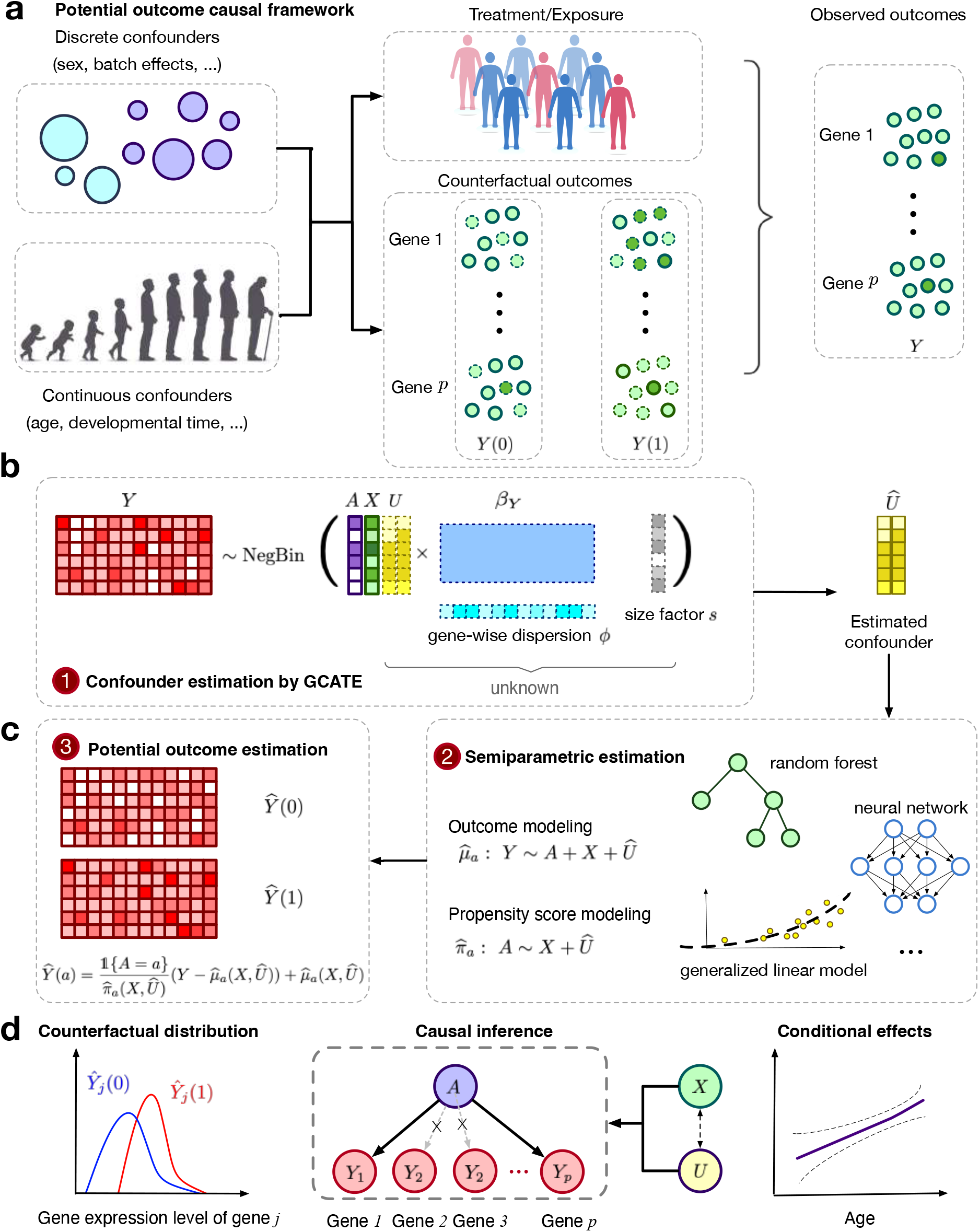
Overview of the proposed causarray method. **a**, Illustration of the data generation process for pseudo-bulk and single-cell data. **b**, In the first step, the gene expression matrix, *Y*, is linked to the treatment, *A*, measured covariates, *X*, and confounding variables, *U*, via a GLM model. The cell-wise size factor, *s*, and gene-wise dispersion parameter, *ϕ*, are estimated from the data, and the unmeasured confounder *U* is estimated by 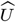 through the augmented GCATE method. **c**, In the second step, generalized linear models and flexible machine learning methods including random forest and neural network can be applied for outcome modeling 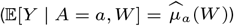 and propensity modeling 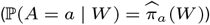 using 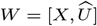. In the third step, the predicted values of the estimated outcome and propensity score functions give rise to the estimated potential outcomes for each cell and each gene. **d**, Downstream analysis includes contrasting the estimated counterfactual distributions, performing causal inference, and estimating the conditional average treatment effects.

We demonstrate the effectiveness of causarray through benchmarking on several simulated datasets, comparing its performance with existing single-cell-level perturbation analysis methods and pseudo-bulk-level differential expression (DE) analysis methods. Next, we apply causarray to two singlecell genomic studies: a Perturb-seq study investigating autism spectrum disorder/neurodevelopmental disorder (ASD/ND) genes in developing mouse brains and a case-control study of Alzheimer’s disease using human brain transcriptomic datasets. For the Alzheimer’s disease analysis, we validate our findings across three independent datasets, showcasing the robustness and reproducibility of causarray in identifying causally affected genes and uncovering biologically meaningful pathways. These applications highlight the potential of causarray to advance our understanding of complex disease mechanisms through rigorous causal inference on general omics.

## Methods

### Counterfactual imputation and inference

Our objective is to determine whether a gene is causally affected by a “treatment” variable after controlling for other technical and biological covariates, which may affect the treatment and outcome variables. Here, we use the term treatment generally; in the narrow sense, it can mean genetic and/or chemical perturbations (26, 27), such as CRISPR-CAS9, and, more broadly, it can mean the phenotype of a disease (7). We acknowledge that while many differentially expressed genes can be considered a result of disease status, for most late-onset disorders, a smaller fraction of genes could have initiated disease phenotypes. Our method aims to determine the direct effects of treatments on modulated gene expression outcomes.

In observational data, the response variable can be confounded by measured and unmeasured biological and technical covariates, making it difficult to separate the treatment effect from other unknown covariates. As a consequence, it is challenging to draw causal inferences; even tests of association may lead to an excess of false discoveries and/or low power. Fortunately, the potential outcomes framework (24, 25) formulates general causal problems in a way that allows for the treatment effect to be separated from the effects of other variables. However, even this framework is challenged by unmeasured covariates. Before introducing our method for estimating unmeasured confounders, we first outline the general potential outcomes framework.

Consider a study in which *Y* is the response variable and *A* is the binary treatment variable for an observation. In the potential outcomes framework, *Y* (*a*) is the outcome that we would have observed if we set the treatment to *A* = *a*. Naturally, we can only observe one of the two potential outcomes for each observation, so

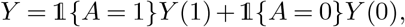

In the context of a case-control study of a disease, this would answer the question: What is the expected difference in gene expression if an individual had the disease (case, *A* = 1) versus if they did not (control, *A* = 0)?

Semiparametric methods provide a powerful tool for estimating potential outcomes in observational studies where randomization is not possible (24, 25). Specifically, we estimate two key quantities: (1) *µ*_*a*_(*X*), the mean response of the outcome variable conditional on treatment *A* = *a* and covariates *X* = *x*, and (2) *π*_*a*_(*X*), the propensity score, which is defined as the probability of receiving treatment *A* = *a* given covariates *X*, i.e., *π*_*a*_(*X*) = ℙ (*A* = *a* | *X*). Using these estimates, we compute potential outcomes using the augmented inverse probability weighted estimator

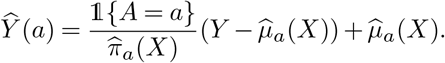

which provides a consistent estimate as long as *either* the outcome model, *µ*_*a*_(*X*), or the propensity score model, *π*_*a*_(*X*), is correctly specified. Given this estimate, we can easily perform downstream inference tasks such as computing log fold change (LFC), and testing for causal effects on gene expressions (Fig. 1a). An advantage of this approach is that counterfactual imputation denoises/balances gene expression under two different conditions. Additionally, having access to estimated potential outcomes facilitates downstream analyses such as estimating causal effects conditional on measured confounders like age.

### The probabilistic modeling of confounders

A key step in these types of analyses is estimating unmeasured confounders. To adjust for confounding, factor models were popularized in surrogate variable analysis literature and have since been widely adopted in bulk gene expression studies (21). We propose an improved version of generalized latent models (18) to identify potential unmeasured confounders, which extends traditional confounder adjustment methods by incorporating more flexible nonlinear models that better capture the unique characteristics of genomic count data, such as zero-inflation (an excess of zero counts) and over-dispersion (greater variability than expected under standard Poisson assumptions). These enhancements allow for more accurate modeling of gene expression data, addressing limitations of simpler linear models in high-dimensional genomic analyses. Using this generalized factor analysis approach, we estimate unmeasured confounders *U* alongside potential outcomes (Fig. 1b-c), enabling direct estimation of downstream quantities such as LFC (Fig. 1d).

We restrict our modelling to one cell type at a time. Although the framework is, in principle, extensible to a joint analysis of all cell types, two practical considerations favour a per-type pipeline: (1) Computational burden. Pooling millions of cells across types inflates both the parameter space and the memory usage, making hyperparameter tuning prohibitively slow for large-scale screens. (2) Dominance of marker-gene signal. When latent factors are learned on the pooled data, the strongest axes of variation are inevitably the marker genes that separate coarse cell identities. As a result, the leading factors reflect cell-type structure rather than the more subtle technical and treatment-related confounders we aim to remove. Preventing this would require ad-hoc marker filtering or explicit constraints, which can be error-prone.

Binning cells by type before estimating confounders, therefore, yields factors that capture within-type heterogeneity (batch, cell-cycle stage, library size, etc.) while leaving biologically meaningful differences between types intact. Whether one ultimately wishes to regress out cell-type variation depends on the scientific objective, but analysing each lineage separately provides a computationally tractable and statistically robust default.

For the *i*th observation (e.g., a single cell or sample) and the *j*th gene, we model the adjusted expression *µ*_*ij*_ = *Y*_*ij*_ */s*_*i*_, where *Y*_*ij*_ is the observed expression level, and *s*_*i*_ is the size factor for the *i*th cell. The size factor accounts for differences in sequencing depth or library size across samples, ensuring that comparisons are not biased by technical variability. We assume that *µ*_*ij*_ follows an exponential family distribution, which is a flexible class of probability distributions commonly used in GLMs with density of *µ*_*ij*_ given by:

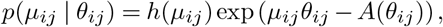

where *θ*_*ij*_ is the natural parameter that determines the mean and variance of *µ*_*ij*_, *h*(*µ*_*ij*_) is a known base measure, and *A*(*θ*_*ij*_) is the log-partition function, which ensures that the density integrates to 1. For negative binomial distributions, both the base measure 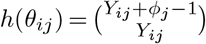 and the log-partition function 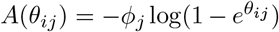 depend on the dispersion parameter *ϕ*_*j*_ of the *j*th gene. The GLMs do not imply that the underlying biological or technical confounders act linearly on expression *Y*_*ij*_. When the dependence between (*A, X*) and the latent confounding is nonlinear, this working model can be viewed as estimating the best low-rank linear projection of that nonlinear nuisance variation in expression space, i.e., the component of unmeasured variation that most strongly explains residual correlation in *Y* after accounting for (*A, X*) and is therefore most relevant for bias reduction.

In matrix form, we model the natural parameters **Θ** = (*θ*_*ij*_)_1≤*i*≤*n*,1≤*j*≤*p*_, as a decomposition into two components: 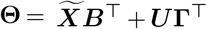. Here, 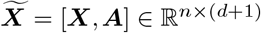 combines observed covariates ***X*** (e.g., biological or technical factors) with treatment indicators ***A***, where *n* is the number of observations, and *d* is the dimension of ***X***; ***B*** ∈ ℝ^*p×*(*d*+1)^ represents unknown regression coefficients for the effects of covariates and treatments on gene expression; ***U*** ∈ ℝ^*n×r*^ represents latent variables capturing unmeasured confounders, where *r* is the number of latent factors; and **Γ** ∈ ℝ^*p×r*^ represents unknown coefficients linking unmeasured confounders to gene expression. This decomposition assumes that gene expression levels are influenced by both observed covariates 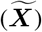 and unmeasured confounders (***U***). The term 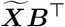 captures the effects of observed covariates and treatments, while ***U* Γ**^⊤^ captures the effects of unmeasured confounders. To estimate the dimension of unmeasured confounders *r* and the corresponding unknown quantities (***B, U***, **Γ**), we employ methods detailed in SI S1.3. This includes techniques for estimating latent factors 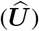 and extending the framework to handle multiple treatments. Once these quantities are estimated, we treat 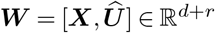 as the complete set of confounding covariates—combining both observed covariates (***X***) and estimated unmeasured confounders 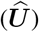.

Single-cell transcriptomic data exhibit systematic variation from both technical and biological sources, even when samples are processed under a nominal “single batch.” Technical effects include library size and capture efficiency (addressed in part via size factors), as well as latent run/lane effects, ambient RNA, and QC-related artifacts. Biological heterogeneity within a cell type (e.g., cell-cycle state, activation/stress programs, and developmental state) also contributes substantial structured variation. Either source can act as a confounder when it influences both the treatment/exposure assignment ***A*** (e.g., disease status or perturbation condition) and gene expression outcomes ***Y***. In causarray, measured technical/biological variables can be included in ***X*** when available, while the remaining shared structure is captured by the estimated latent factors 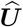 under the generalized factor model 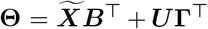, yielding 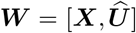 for downstream semiparametric causal estimation.

### Semiparametric estimation

Throughout the paper, we consider the LFC as the target estimand

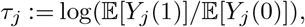

which quantifies the relative change in expected gene expression levels between treatment (*A* = 1) and control (*A* = 0) conditions for gene *j*. Extensions to other estimands are provided in SI S2.2. The semiparametric estimation framework is a widely used approach that is agnostic to the underlying data-generating process. It provides valid estimation and inference results as long as either the conditional mean model (*µ*_*j*_) or the propensity score model (*π*) is correctly specified. This robustness property ensures reliable causal effect estimation even in the presence of potential misspecification of one of the models.

More specifically, a one-step estimator 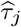 of the estimand *τ*_*j*_ admits a linear expansion:

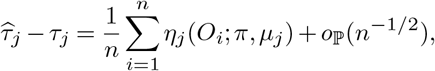

where *η*_*j*_(*O*_*i*_; *π, µ*_*j*_) is the influence function of *τ*_*j*_, which quantifies how individual observations contribute to the overall estimate. Here, *π*(***W***) = ℙ (*A* = *a* | ***W***) is the propensity score model, and *µ*_*j*_(***W***, *a*) = 𝔼 [*Y*_*j*_ | *W, A* = *a*] is the outcome model for gene *j*.

To estimate the nuisance functions *µ*_*j*_’s (outcome models) and *π* (propensity score model), we use flexible statistical machine learning methods. Specifically, for outcome models *µ*_*j*_, we employ generalized linear models (GLMs) with a negative binomial likelihood and log link function. This choice accounts for over-dispersion in count data while ensuring computational efficiency given the high dimensionality of genomic data. For the propensity score model *π*, we provide two built-in options: (i) logistic regression and (ii) random forests. In our experiments with Alzheimer’s disease analysis with mixed-type covariates, the random forest model is configured for robustness and performance. Default parameters for each tree include a minimum of 10 samples per leaf, a minimum of 20 samples required to split a node, and the number of features considered for each split set to the square root of the total number of features (max features=‘sqrt’). To account for imbalanced treatment groups, class weights are automatically balanced. Other hyperparameters are then tuned using Extrapolated cross-validation (ECV) (28) to improve the Gini impurity. Specifically, the number of trees from 200 to 1000, the maximum tree depth is selected from {3, 5, 7} and the proportion of samples used to train each tree (max_samples) is selected from {0.4, 0.6, 0.8, 1.0}. Minimal cost-complexity pruning is also applied with a ‘ccp alpha’ of 0.02. Users can also supply alternative estimates for these nuisance functions if desired.

To perform inference, we first compute the estimated influence function values 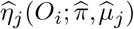 and use them to estimate the variance for gene *j*:

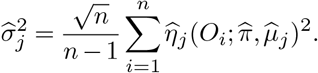

Using these quantities, a two-sided *t*-statistic for gene *j* can be computed as: 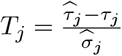. This statistic enables hypothesis testing and confidence interval construction for causal effects on gene expression. For further details on the method, see SI S1-S2.

### Computational considerations

The computational cost of causarray is driven by two components: (i) fitting the generalized factor model to estimate *U*, and (ii) semiparametric effect estimation and inference across genes. Both components are highly parallelizable across genes, and the memory footprint is dominated by storing the count matrix and low-rank factors.

## Results

### Simulation study demonstrates the advantages of causarray

We evaluate the performance of causarray in two simulated settings (SI S3). In the first setting, we generate simulated pseudo-bulk data, while in the second, we generate simulated single-cell data using the Splatter simulator (29), which explicitly models the hierarchical Gamma-Poisson processes underlying scRNA-seq data and captures multi-faceted variability. Each dataset consists of 100-5000 pseudo-bulk or single-cell observations, approximately 2,000 genes, 1-2 covariates, and 4 unmeasured confounders.

To benchmark causarray, we compare it with several existing methods designed for differential expression (DE) testing, both with and without confounder adjustment (Fig. 2a). For methods that do not account for unmeasured confounders, we include the Wilcoxon rank-sum test and DESeq2 (30). In the presence of measured covariates, both regress the gene expression counts with respect to the covariates using the Poisson or negative binomial generalized linear model, respectively. The input to the Wilcoxon rank sum test is the deviance residuals. For confounder-adjusted methods, we consider CoCoA-diff (7), CINEMA-OT (20), CINEMA-OTW (20), Mixscape (31), RUV (22), and RUV-III-NB (32), where recommended DE test methods are subsequently applied with estimated confounders. RUV estimates latent confounders using RUVSeq (22), followed by DESeq2 (30) for differential testing. RUV-III-NB extends RUVSeq by modeling count data directly using a negative binomial distribution, which improves robustness in overdispersed single-cell RNA-seq data. Mixscape reduces confounding from treatment heterogeneity by using matching to identify and isolate truly responding cells before comparing them to controls for differential expression analysis. CINEMA-OT employs a heuristic independent space analysis procedure plus optimal transport to separate confounding variables from treatment-associated variables. Compared to CINEMA-OT, CINEMA-OT-W further adjusts for differences in propensity scores between treated and control cells by matching before independent component analysis. A brief summary of each benchmarking comparison method is provided in SI S3.2.

**Fig. 2.**
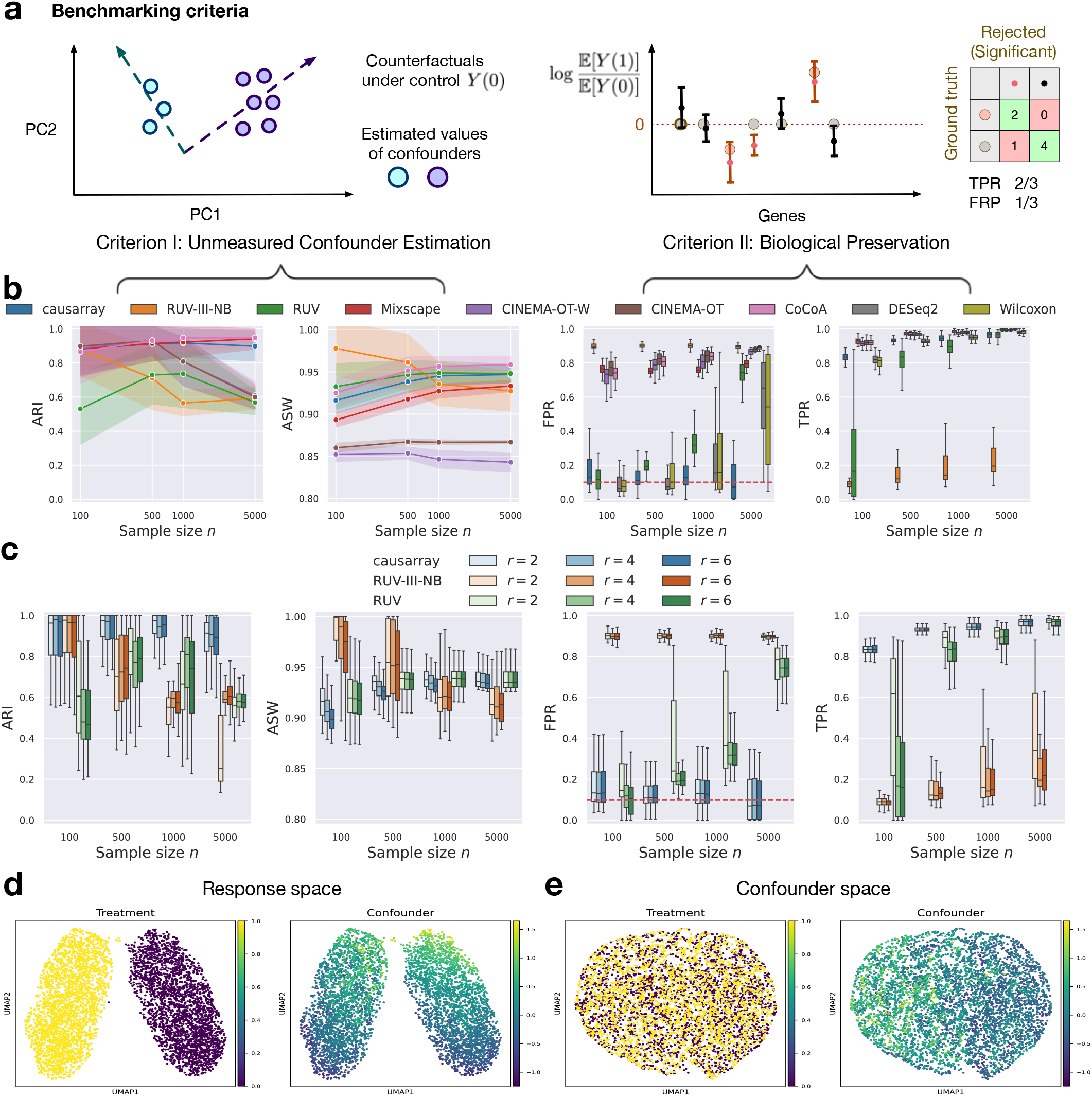
Benchmarking of causarray against other methods for single-cell differential expression testing on synthetic expression data with unmeasured confounders. **a**, The analysis pipeline produces a confounder adjustment and a statistic for DE testing. We illustrate two types of criteria used for benchmarking confounder adjustment and DE methods in simulations for pseudo-bulk expressions (**b-e**) and single-cell (Fig. S2) expressions. The metrics for each experimental setting are calculated using 50 simulated datasets with varying random seeds. **b**, Performance comparison of causarray and other methods with a well-specified number of latent factors (*r* = 4). Line plots show mean ARI and ASW scores for confounder estimation (shaded region represents values within one standard deviation), while box plots display FPR and TPR for biological signal preservation. For the box plots, the center indicates the median, the top and bottom hinges represent the top and bottom quartiles, and the whiskers extend from the hinge to the largest or smallest value no further than 1.5 times the interquartile range from the hinge. **c**, Robustness analysis of causarray, RUV-III-NB, and RUV under varying numbers of latent factors (*r* = 2, 4, 6). **d-e**, causarray disentangles the treatment effects and unmeasured confounding effects in the response and confounder spaces (with *n* = 5000 samples). UMAP projection of (**d**) expression data *Y* colored by the values of treatment *A* (purple for control *A* = 0 and yellow for treated *A* = 1) and unmeasured continuous confounder *U*; and (**e**) estimated potential outcome under control *Y* (0) colored by the values of treatment *A* and continuous confounder *U*.

To assess the performance of unmeasured confounder adjustment procedures, we use two metrics: adjusted Rand index (ARI) and average silhouette width (ASW). More specifically, we use ARI to quantify the alignment between estimated and true unmeasured confounders and ASW to evaluate cell type separation in the control response space. A higher ARI value indicates better coherence, and a higher ASW value reflects better preservation of biological signals after removing confounding effects. Additionally, to assess the performance of DE testing, we use two metrics: false positive rate (FPR) and true positive rate (TPR) (SI S3.3).

We first evaluate how sample size and confounding levels influence the performance of DE testing across methods. Among all tested approaches, only causarray, RUV, Wilcoxon, and DESeq2 effectively control FPR with moderate sample sizes (*n* ≤ 500) (Fig. 2b and Fig. S2ab). Once the sample size grows larger (*n>* 500), only causarray continues to maintain FPR close to the nominal level of 0.1 across all confounding levels, whereas the FPR of RUV, Wilcoxon, and DESeq2 drifts upward to ~0.15–0.22. On the other hand, Mixscape, CINEMA-OT, CINEMA-OT-W, RUV-III-NB, and CoCoA-diff exhibit inflated FPRs exceeding 0.5 in most cases. Within the subset of methods that control FPR reasonably well (causarray, RUV, Wilcoxon, DESeq2), causarray achieves the highest TPRs, consistently reaching 0.80–0.90 across all scenarios (Fig. 2b). This is significantly higher than DESeq2 and Wilcoxon, particularly for smaller sample sizes in the single-cell experiments (Fig. S2ab). These results highlight causarray’s ability to balance sensitivity and specificity effectively.

In terms of unmeasured confounder adjustment, causarray, RUV-III-NB, and CoCoA-diff achieve both ARI and ASW scores consistently above 0.7 across all sample sizes in both pseudo-bulk and single-cell data (Fig. 2b, Fig. S2ab), outperforming RUV in ARI and CINEMA-OT-W/CINEMA-OT in ASW. Furthermore, causarray effectively disentangles treatment effects from unmeasured confounding effects. In the response space (Fig. 2d), treatment groups are distinctly separated with minimal overlap, while variations within groups reflect unmeasured confounders. In the confounder space (Fig. 2e and Fig. S1), causarray produces a uniform mixing of treatment groups while accurately reconstructing continuous confounder values.

Finally, we assess the robustness of causarray, RUV-III-NB, and RUV under varying numbers of latent factors (Fig. 2c and Fig. S2c). Among these methods, only causarray consistently controls FPR at nominal levels of 0.1, regardless of the number of factors or sample size. In contrast, RUV-III-NB exhibits inflated median FPRs exceeding 0.2 when more factors are included (e.g., *r* = 6). While RUV-III-NB performs well in terms of ARI (above 0.8) and ASW (above 0.7), its DE testing performance is inferior to RUV due to poor FPR control under certain conditions. Based on these findings, we proceed with causarray and RUV for real data analysis.

### An in vivo Perturb-seq study

#### An integrative analysis of multiple single perturbations

Autism spectrum disorders and neurodevelopmental delay (ASD/ND) represent a complex group of conditions that have been extensively studied using genetic approaches. To investigate the underlying mechanisms of these disorders, researchers have employed scalable genetic screening with CRISPR-Cas9 technology (26). Frameshift mutations were introduced in the developing mouse neocortex in utero, followed by single-cell transcriptomic analysis of perturbed cells from the early postnatal brain (26). These in vivo single-cell Perturb-seq data allow for the investigation of causal effects of a panel of ASD/ND risk genes. We analyze the transcriptome of cortical projection neurons (excitatory neurons) perturbed by one risk gene or a non-targeting control perturbation, which serves as a negative control (see SI S3.4 for details).

Unmeasured confounders, such as batch effects and unwanted variation, are likely present in this dataset due to the batch design being highly correlated with perturbation conditions (Fig. S3ab). Additionally, the heterogeneity of single cells assessed in vivo introduces further complexity. These confounding factors may reduce statistical power for gene-level differential expression (DE) tests, as noted in the original study (26), which instead focused on gene module-level effects. To address this limitation, we apply causarray to incorporate unmeasured confounder adjustment and conduct a more granular analysis at the single-gene level. This approach enables us to uncover nuanced genetic interactions and causal effects that may provide deeper insights into the etiology of ASD/ND.

#### Functional analysis

Gene module-level analyses have been shown to provide greater statistical power for detecting biologically meaningful perturbation effects when fewer cells are available (26). The original study adopted this approach but relied on a linear model rather than a negative binomial model, potentially limiting its ability to detect broader signals at the individual gene level. Here, we compare causarray with RUV and DESeq2 (without confounder adjustment) to identify significant genes and enriched gene ontology (GO) terms associated with various perturbations. The number of latent factors is set as 10, according to the joint-likelihood-based information criterion (SI S1.3, Fig. S4a).

In terms of significant gene detection, causarray identifies a comparable number of significant genes to RUV across most perturbations, while DESeq2 consistently detects fewer significant genes (Fig. 3a). The variation in significant detections across different perturbed genes suggests distinct biological impacts of each knockout. Functional analysis focuses on enriched GO terms on the DE genes under each perturbation condition where discrepancies arise between causarray and other methods. Genes identified by causarray are enriched for biologically relevant GO terms with clear clustering patterns (Fig. 3b-c, Fig. S3c). In contrast, RUV shows less distinct clustering and enrichment patterns.

**Fig. 3.**
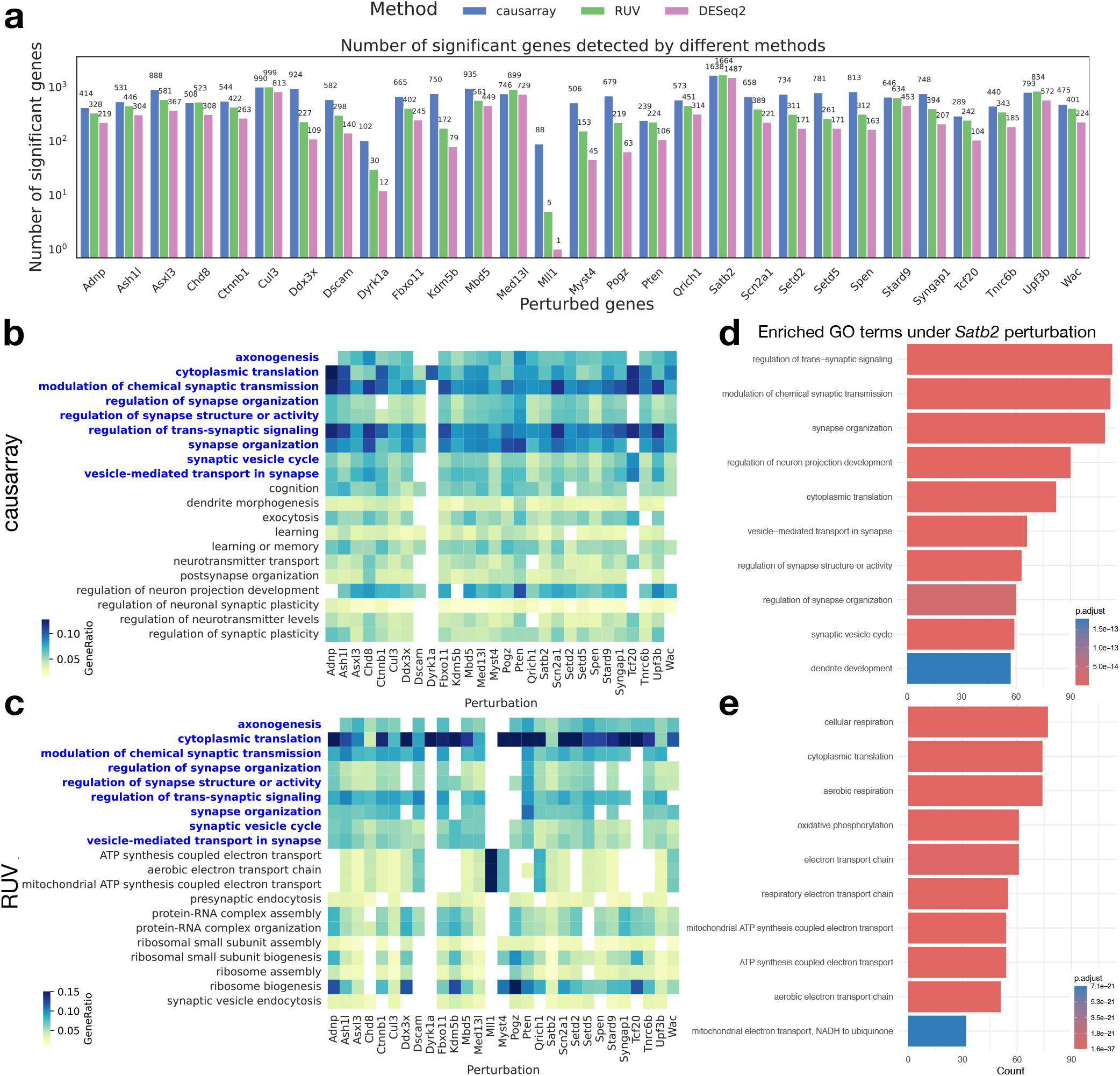
Statistical test results of the effects of CRISPR perturbation on gene expression in excitatory neuron data. **a**, Number of significant genes detected under all perturbations using three different methods. The detection threshold for significant genes is FDR*<* 0.1 for all methods. **b-c**, Heatmaps of GO terms enriched (adjusted *P* value *<* 0.05, *q <* 0.2) in discoveries from causarray and RUV, respectively, where the common GO terms are highlighted in blue. Only the top 20 GO terms that have the most occurrences in all perturbations are displayed. **d-e**, Barplots of GO terms enriched in discoveries under *Satb2* perturbation from causarray and RUV, respectively.

Notably, while RUV identifies GO terms related to ribosome processes previously implicated in ASD studies (33), these findings remain controversial. Some argue that dysregulation in translation processes and ribosomal proteins may reflect secondary changes triggered by expression alterations in synaptic genes rather than direct causal effects (34). In contrast, GO terms identified by causarray align more closely with the expected causal effects of ASD/ND gene perturbations (35, 36).

To further validate these findings, we examine the perturbation condition for *Satb2*, which yields the largest number of significant genes identified by both methods (adjusted *P* value *<* 0.1) and exhibits significantly different estimated propensity scores (Fig. S4b). *Satb2* is known to play critical roles in neuronal development, synaptic function, and cognitive processes (37, 38). Using causarray, we detect enrichment for GO terms directly related to neuronal function and development, such as “regulation of neuron projection development,” “regulation of synapse structure or activity,” and “synapse organization” (Fig. 3d). These findings are consistent with *Satb2*’s established roles in neuronal development and synaptic plasticity (39, 40). On the other hand, RUV identifies enrichment for terms related to mitochondrial function and energy metabolism, such as “mitochondrial electron transport,” “cellular respiration,” and “ATP synthesis” (Fig. 3e). While these processes are important for general cellular function, they are less directly relevant to *Satb2*’s primary biological roles. For an extended list of GO terms, see Fig. S3c.

A gene-level breakdown under *Satb2* (including causarrayonly genes in GO:0021953 and their association with the estimated latent factors 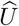) is provided in SI S3.4 (Fig. S5ab). We additionally applied ComBat (41) as a representative design-based correction method that requires known batch labels. In the *Satb2* perturbation, ComBat identifies 1710 DE genes, but yields no GO biological process terms passing multiple-testing correction (adjusted *P <* 0.05), whereas causarray identifies 1638 DE genes and RUV identifies 1705 (FDR *<* 0.1) (Fig. S5cd). Moreover, ComBat’s top GO terms are broad, high-level categories (e.g., signaling, phosphorylation, cell adhesion), while *Satb2* knockdown is expected to enrich neuronal development and synaptic programs. This illustrates a practical failure mode when batch design is strongly aligned with perturbation assignment: design-based correction can reduce biological specificity, motivating causal estimation procedures that explicitly model treatment assignment and latent confounding.

Overall, this analysis demonstrates that causarray provides greater specificity in detecting biologically meaningful causal effects of gene perturbations. Its ability to disentangle confounding influences while preserving relevant biological signals highlights its effectiveness in analyzing complex genomic datasets.

### Alzheimer’s disease case-control study

#### An integrative analysis of excitatory neurons

We analyze three Alzheimer’s disease (AD) single-nucleus RNA sequencing (snRNA-seq) datasets: a transcriptomic atlas from the Religious Orders Study and Memory and Aging Project (ROSMAP) (42) and two datasets from the Seattle Alzheimer’s Disease Brain Cell Atlas (SEA-AD) consortium (43), which include samples from the middle temporal gyrus (MTG) and prefrontal cortex (PFC). Our objective is to compare the performance of causarray and RUV in pseudo-bulk DE tests of AD in excitatory neurons (see SI S3.5 for details).

To evaluate the validity, we perform a permutation experiment on the ROSMAP-AD dataset by permuting phenotypic labels. Ideally, no significant discoveries should be made under this null scenario. However, RUV produces a large number of false discoveries, with its performance deteriorating as the number of latent factors increases. In contrast, causarray effectively controls the false discovery rate (FDR), producing minimal false positives (Fig. 4a), and the results using FDX are conservative. Additionally, we assess coherence across datasets by examining effect sizes in SEA-AD (MTG) and SEA-AD (PFC). Because we cannot know the ground truth, our objective is to assess the stability of results across the three related studies. Effect sizes estimated by causarray exhibit higher consistency across varying q-value cutoffs compared to RUV (Fig. 4bc). We further compare functional enrichment results between causarray and RUV using gene ontology (GO). When inspecting DE genes across all three AD datasets, causarray identifies similar numbers of discoveries as RUV, and 1186 terms were associated with them (SI S3.5, Fig. S6b). The discovered networks, as defined as the top 5 GO terms and associated genes included in the top 100 DE gene discoveries, show the enhanced sensitivity and comprehensiveness of causarray (Fig. 4d).

**Fig. 4.**
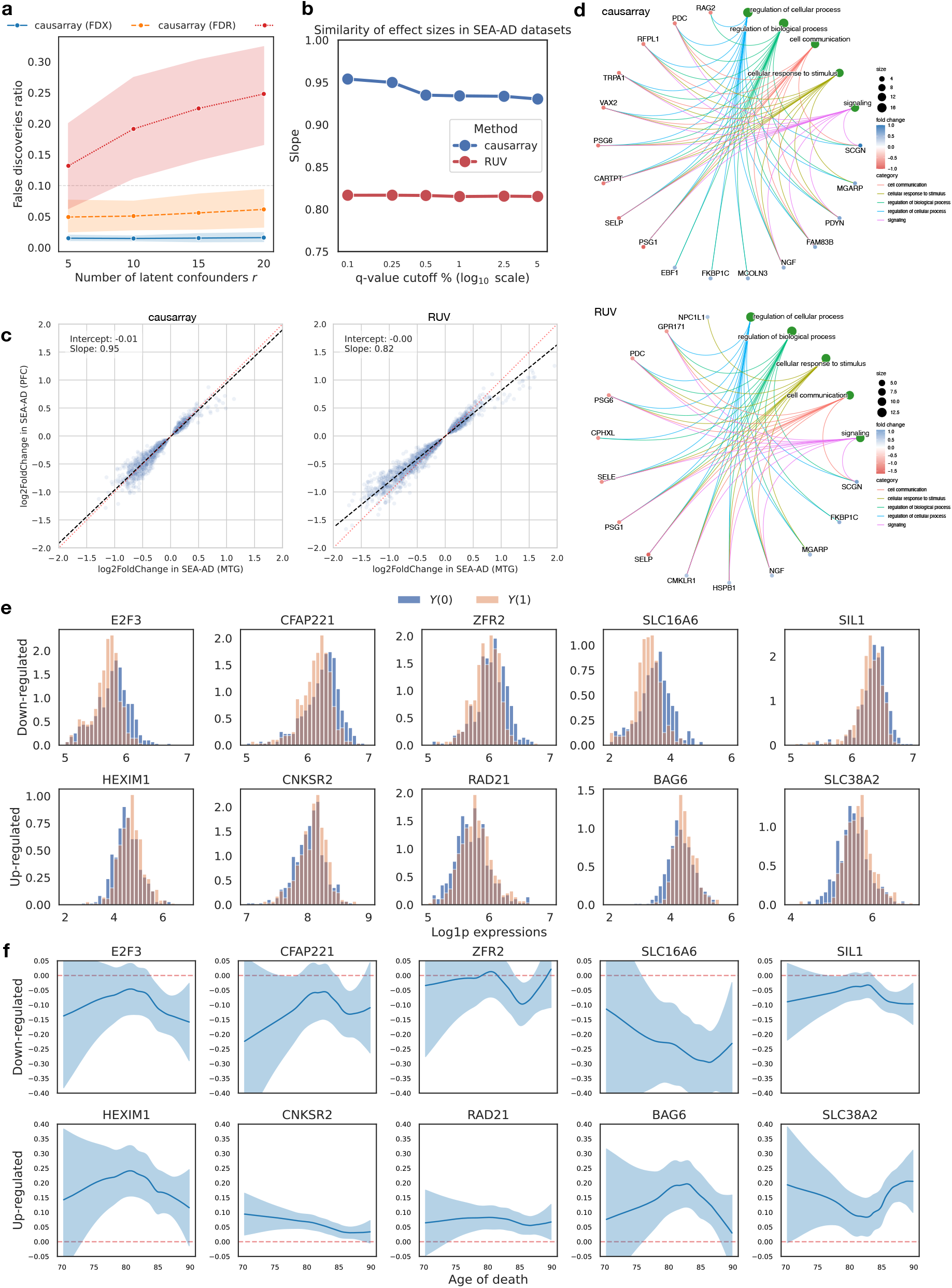
Comparison of DE genes discovered by causarray and RUV on excitatory neurons for Alzheimer’s disease. **a**, The ratio of false discoveries to all 15586 genes of DE test results with permuted disease labels on the ROSMAP-AD dataset. Three methods, causarray with FDX control, causarray with FDR control, and RUV with FDR control, are compared. Data are presented as mean values *±* s.d. **b**, The similarity of estimated effect sizes on SEA-AD MTG and PFC datasets. The slope is estimated from linear regression of effect sizes on the PFC dataset against those on the MTG dataset. **d**, Considering only the top 50 positively regulated and the top 50 negatively regulated DE genes from causarray and RUV, we map them to the top 5 biological processes (the green nodes). **e**, Estimated counterfactual distributions of 10 selected genes by causarray. The top 5 up-regulated and top 5 down-regulated genes in estimated LFCs (adjusted *P* value *<* 0.05) are visualized. The values are displayed on a log scale after adding a pseudo-count of one. **f**, Estimated log-fold change of treatment effects, conditional on age for selected genes. The center lines represent the mean of the locally estimated scatter plot smoothing (LOESS) regression, and the shaded area represents a 95% confidence interval at each value of age.

#### Counterfactual analysis

The counterfactual framework employed by causarray enables downstream analyses that directly utilize estimated potential outcomes. By examining counterfactual distributions for significant genes (Fig. 4a), we observe distinct shifts in expression levels between treatment (*Y* (1)) and control (*Y* (0)) groups. Downregulated genes show a shift toward lower expression levels under disease conditions, while upregulated genes exhibit increased expression. Conditional average treatment effects (CATEs) reveal age-dependent trends for these genes (Fig. 4b). For example, upregulated genes such as *SLC16A6* and *HEXIM1* show stronger effects at extreme ends of the age distribution, while others like *SLC38A2* and *BAG6* display nuanced changes across the aging spectrum.

These findings align with prior studies highlighting the roles of specific genes in aging-related processes. For instance, *ZFR2, SIL1, BAG6*, and *RAD21* have been implicated in chromatin remodeling, synaptic plasticity, and cellular stress responses critical for aging and neurodegeneration (44–47). While nonparametric fitted curves exhibit wider uncertainty bands, particularly at the boundaries, which can be observed here, the significant trends observed for key genes highlight their potential relevance in AD pathology. Overall, these results demonstrate that causarray provides nuanced insights into age-dependent gene regulation mechanisms while maintaining robust control over confounding influences.

## Conclusion

The rapid growth of high-throughput single-cell technologies has created an urgent need for robust causal inference frameworks capable of disentangling treatment effects from confounding influences. Existing methods, such as CINEMAOT (20), have advanced the field by separating confounder and treatment signals and providing per-cell treatment-effect estimates. However, these methods rely on the assumption of no unmeasured confounders, which is often violated in observational studies and in vivo experiments. Additionally, many confounder adjustment methods, such as RUV (22), depend on linear model assumptions that do not directly model count data or provide robust differential expression testing at the gene level. Similarly, popular single-cell integration tools primarily target cross-batch alignment in a transformed or embedded space rather than producing corrected count-scale expression for DE testing (SI S1.6). Likewise, while DESeq2 (30) is a widely used tool for differential expression analysis of count data, it assumes that treatment assignment is unconfounded after accounting for observed covariates and does not explicitly model latent confounding. In contrast, causarray builds on the generalized linear modeling framework to estimate unmeasured confounders and extend the standard parametric models to a semiparametric regime with flexible nuisance function estimation. This distinction is critical in single-cell and case–control studies where hidden batch or biological effects can otherwise bias DESeq2’s inference.

causarray directly models count data using generalized linear models for unmeasured confounder estimation, overcoming a key limitation of RUV in DE analysis. Unlike CINEMAOT (20) and CoCoA-diff (7), which rely on optimal transport or matching techniques, causarray employs a semiparametric framework that combines flexible machine learning models with semiparametric inference. This approach enhances stability and interpretability while enabling valid statistical inference of treatment effects. Benchmarking results demonstrate that causarray outperforms existing methods in disentangling treatment effects from confounding influences across diverse experimental settings, maintaining superior control over false positive rates while achieving higher true positive rates. The use of generalized linear models allows more accurate estimation of latent confounders in sparse, overdispersed count data, improving the robustness of confounder adjustment relative to linear or heuristic matching approaches. By explicitly modeling the mean–variance relationship of single-cell data, the GLM formulation helps extract biologically relevant latent structure that would otherwise be absorbed as noise. Nevertheless, errors in the estimated confounders 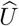 can propagate into downstream causal estimates, leading to bias or inflated variance when the latent structure is weakly identifiable. Accordingly, causal interpretation in causarray is conditional on standard identification assumptions (consistency, conditional exchangeability given (*X*, 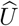), and positivity) and on the adequacy of the estimated latent structure for capturing residual shared variation relevant to treatment assignment and expression. When overlap is limited, or the latent structure is weakly identifiable, estimates can be sensitive to modeling choices (including *r*) and should be interpreted as robustness-focused causal evidence under assumptions rather than as unconditional causal ground truth. These effects are inherent to all confounder-adjustment approaches and underscore the importance of diagnostic checks, such as overlap assessment and sensitivity analyses, to evaluate the robustness of causal conclusions.

In an in vivo Perturb-seq study of ASD/ND genes, causarray uncovered gene-level perturbation effects that were missed by prior module-based analyses. The Perturb-seq experiment involved 30 distinct perturbation conditions, all analyzed jointly by causarray within a unified framework that adjusts for shared confounding across groups. It identified biologically relevant pathways linked to neuronal development and synaptic functions for multiple autism risk genes. Similarly, in a case-control study of Alzheimer’s disease using three human brain transcriptomic datasets, causarray revealed consistent causal gene expression changes across datasets and highlighted key biological processes such as synaptic signaling and cell development. The same framework is readily extensible to multi-group comparisons in disease studies—for example, different stages of Alzheimer’s progression—allowing stage-specific effect estimation while accounting for shared sources of variation. These findings underscore the ability of causarray to provide biologically meaningful insights across diverse contexts.

Despite its strengths, causarray has certain limitations. Its performance depends on the accurate estimation of unmeasured confounders, which may vary with dataset complexity and experimental design. Furthermore, while causarray provides robust DE testing, its integration with advanced spatial or trajectory analysis frameworks remains unexplored (48, 49). Future research could focus on extending causarray to incorporate prior biological knowledge or extrapolate to unseen perturbation-cell pairs, similar to emerging methods like CPA (50). Such advancements would further enhance its applicability in single-cell causal inference on general omics.

Finally, in this study, we focus our analysis on scRNA-seq outcomes, but emerging technologies now enable a wide range of omic readouts. In the perturbation framework, these include chromatin-based readouts with Perturb-ATAC (51), protein-level profiling with ProCODE (52), and joint RNA–protein measurements with ECCITE-seq (53). causarray can easily be extended to any of these readouts.

## KEY POINTS

- We introduce causarray, a causal inference framework for single-cell and pseudo-bulk omics that couples generalized confounder adjustment for count data with semiparametric inference.
- In simulations, causarray disentangles treatment effects from unmeasured confounding, maintains nominal false-positive control, and achieves higher power while preserving biological structure across sample sizes and latent-factor choices.
- In a benchmarking study, causarray performs better than other methods, both in terms of confounder estimation and testing performance (true positives and true negatives).
- Applied to in vivo Perturb-seq of ASD/ND genes, causarray detects gene-level causal effects enriched for neuronal development and synaptic pathways, offering greater specificity than alternative methods.
- In Alzheimer’s case-control analyses spanning ROSMAP and SEA-AD cohorts, causarray yields reproducible effect sizes, rigorous error-rate control, and clear counterfactual shifts in expression that highlight synaptic and cell-development processes.
- By estimating per-gene potential outcomes, causarray enables downstream counterfactual and conditional treatment-effect analyses (for example, age-dependent trends), providing a practical route from association toward causal effect estimation under explicit identification assumptions.

## CODE AVAILABILITY

The code for reproducing the results in the paper and the causarray package can be accessed at https://github.com/jaydu1/causarray.

## DATA AVAILABILITY

All datasets used in this paper are previously published and freely available, except the metadata for donors from the ROSMAP cohort. The Perturb-seq dataset is available through the [Broad single cell portal](https://singlecell.broadinstitute.org/single_cell/study/SCP1184) as txt files. The gene expression count matrices of ROSMAP-AD datasets (42) can be obtained from [supplementary website](https://compbio.mit.edu/ad_aging_brain/#processed-count-matrices), which have been deidentified to protect confidentiality - the mapping to ROSMAP IDs and complete metadata can be found on [Synapse](https://compbio.mit.edu/ad_aging_brain/#processed-count-matrices) as Seurat objects (rds files). The SEA-AD datasets of nuclei-by-gene matrices with counts and normalized expression values from the snRNA-seq assay (43) are available through the [Open Data Registry](https://registry.opendata.aws/allen-sea-ad-atlas/) in an AWS bucket (sea-ad-single-cell-profiling) as AnnData objects (h5ad files).

## ACKNOWLEDGEMENTS

The first author gratefully acknowledges support from the Institute for Data Science (IDS) at the University of Hong Kong through the HKU100 award. Part of the computations was performed using research computing facilities offered by Information Technology Services, the University of Hong Kong, and the Bridges-2 system at the Pittsburgh Supercomputing Center (PSC) through allocation MTH230011P from the Advanced Cyberinfrastructure Coordination Ecosystem: Services & Support (ACCESS) program. This project was funded by the National Institute of Mental Health (NIMH) grant R01MH123184.

## Supplementary Information S1: Confounder estimation

### S1.1 Unit of analysis and dependence

When many cells are sampled from the same donor/subject, within-donor correlation violates an i.i.d. cell-level assumption; in such settings, we recommend donor-level pseudo-bulk aggregation and treat donors as the experimental units (as in our Alzheimer’s disease analyses). Cell-level analyses are most appropriate when cells are approximately independent experimental units (e.g., cell lines or weakly clustered designs).

### S1.2 Algorithm

To estimate the unmeasured confounders, we employ an improved version of GCATE (18). Suppose (*X*_*i*_, *A*_*i*_, *Y*_*i*_) for *i* = 1, …, *n* are *n* independently and identically distributed samples coming from the same distribution as (*X, A, Y*) ∈ ℝ^*d*^ *×* ℝ^*a*^ *×* ℝ^*p*^. Here, *A* consists of *a* treatments and can be both continuous and discrete for the purpose of confounder estimation. Let ***X*** ∈ ℝ^*n×d*^, ***A*** ∈ ℝ^*n×a*^, ***Y*** ∈ ℝ^*n×p*^ denote the design matrix, treatment matrix, and gene expression matrix, respectively. To account for different library sizes, we model the mean of the size-normalized counts

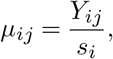

which is assumed to follow a negative binomial distribution. Technically, *µ*_*ij*_ ‘s should be non-negative integers; however, the likelihood-based approaches work seamlessly even when they are non-negative real numbers. Here *s*_*i*_ is the size factor of cell *i*, which will be specified later. We assume the conditional mean is characterized by a generalized linear model

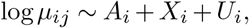

and its dispersion parameter *ϕ* is predetermined.

The adjusted expression *µ*_*ij*_ of the *i*th observation and the *j*th gene has the density:

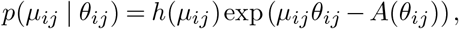

where *θ*_*ij*_ is the natural parameter. In matrix form, the natural parameters decompose as

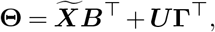

Where 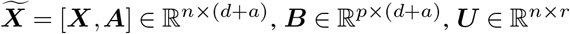, and **Γ** ∈ ℝ^*p×r*^ are unknown. Note that *µ*_*ij*_’s are conditionally independent given the natural parameter **Θ**. Under such a model, the theoretical foundations for identifying *U* are provided in prior work (18), which establishes conditions under which ***U*** can be consistently estimated from observed data using matrix factorization techniques, even in the presence of latent confounding. With this notation, the procedure of unmeasured confounder estimation is summarized in Algorithm S1, and the details of the method are described below.

#### Estimation of size factors

We follow the procedure in (30) to compute the size factors *s*_*i*_ for *i* = 1, …, *n*. We start by calculating the geometric mean for each gene *j*:

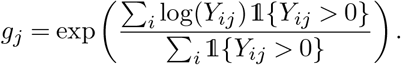

Next, for each sample *i*, compute the initial size factors:

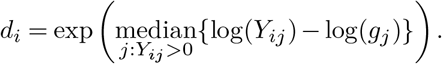

Finally, we normalize these size factors to have a geometric mean of 1 across all samples:

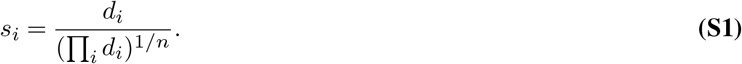

The size factors can then be used to normalize gene expression data, adjusting for differences in sequencing depth and other systematic biases across samples. The normalization ensures that observed differences in expression levels reflect true biological variation rather than technical artifacts.

##### Algorithm S1

Unmeasured confounder estimation

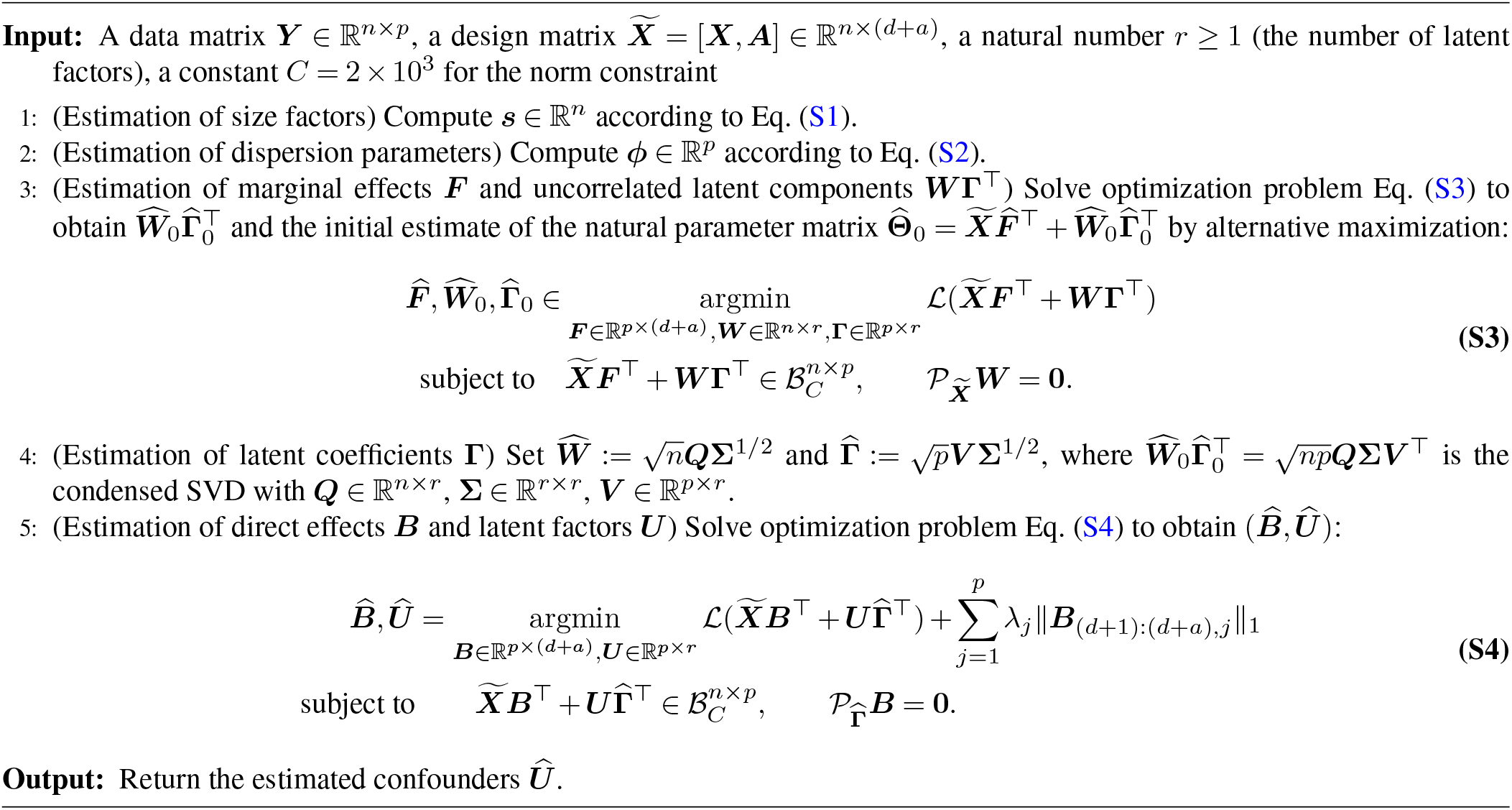

#### Estimation of dispersion parameters

To estimate the dispersion parameter, we first fit generalized linear models (GLMs) on the data and obtain the estimated mean expression of gene *j*, denoted as 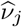 for *j* = 1,…, *p*. Note that when *µ*_*ij*_ comes from a Negative Binomial distribution, its variance is given by

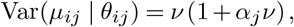

where *ν* = 𝔼 [*µ*_*ij*_ | *θ*_*ij*_] is the conditional mean while *α*_*j*_ is the dispersion parameter of the NB1 form. In the form of an exponential family parameterized by the parameter *ϕ*_*j*_, *α*_*j*_ is the reciprocal of *ϕ*_*j*_, namely, *α*_*j*_ = 1*/ϕ*_*j*_. By methods of moments, we can solve the following equation to obtain an estimator 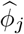 for *ϕ*_*j*_:

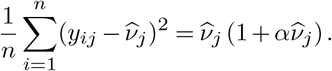

Finally, we clip 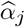 to be in [10^−2^, 10^2^] and set 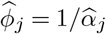. The estimated dispersion parameter has a closed-form expression:

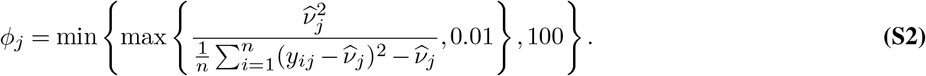

#### Estimation of marginal effects by joint likelihood estimation

The negative log-likelihood function of the data is given by

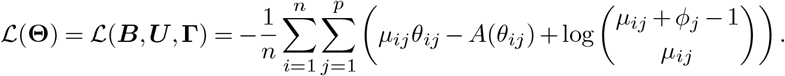

Although this is a nonconvex optimization problem, an alternative descent algorithm as in (18) can be employed to solve it efficiently. By rewriting 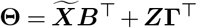 as 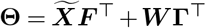 with 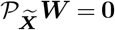, we can disentangle the marginal effects and the uncorrelated latent components. This corresponds to step 3 of Algorithm S1. Each entry of the estimated natural parameter matrix is constrained within the Euclidean ball ℬ_*C*_ with radius *C* (*C* = 2 *×* 10^3^ by default).

Before alternative maximization, we compute deviance residuals ***R*** from the NB GLM fits with offsets log ***s*** and dispersion parameters *ϕ*, and initialize the uncorrelated confounders by 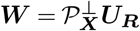 where ***U***_***R***_ ∈ ℝ^*n×r*^ contains the first *r* left singular vectors of ***R***. Here, the projection 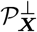 ensures that ***W*** is uncorrelated with ***X***. Then, we initialize the marginal effects ***F*** and latent coefficient **Γ** by solving GLMs with covariates 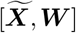. In particular, when the intercept is included in the covariates, the initial value of ***W*** also has zero means per column.

#### Estimation of latent coefficients

Because the (uncorrelated) latent factors are identifiable only up to scaling and rotations, we rescle the estimate at step 4 of Algorithm S1. This ensures the eigenvalues of 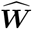 and 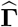 have the same order, making the alternative optimization more stable.

#### Estimation of confounding effects by adaptive penalization

The last step is to jointly recover the direct effects and the unmeasured confounders. This is done by imposing orthogonality between 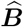 and 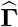, as well as imposing sparsity on 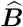. The former ensures the gene-wise effects of the observed covariates and the unmeasured confounders are uncorrelated, while the latter aims to reveal signals from noisy measurements.

The direct effect ***B*** is initialized as 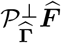. Then, Initialize ***Z*** and **Γ** using the SVD of the matrix 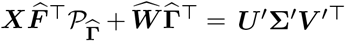. Let ***Z*** = (***V*** ′**Σ**′^1/2^)_1:*r*_ and ***Z*** = (***V*** ′**Σ**′^1/2^)_1:*r*_ be the initialized values.

To account for different scales of the effects induced by different treatment conditions, we propose to use the adaptive lasso to induce sparsity of effects from multiple treatments. The coefficients for non-treatment covariates ***X*** are not penalized in our implementation. Only the treatment-related coefficients for ***A*** are subject to penalization. More specifically, in optimization problem S4, the regularization parameters are set as

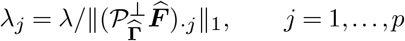

where 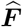 is the design matrix and 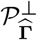 projects out estimated confounding effects. This scaling adjusts for the varying magnitudes across genes and ensures that the penalization is adaptive to gene-specific signal strength, in contrast to using a single global *λ* as in GCATE (18). In our implementation, we use a default value of *λ* = 0.05, which was chosen based on empirical performance across several datasets. Results are not overly sensitive to small variations in *λ* due to the gene-wise normalization. Users can also tune *λ* via cross-validation if needed.

Because of regularization, the estimate 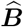 is typically biased towards zero, so we don’t use it for downstream analysis. It is possible to perform inference with an additional debiasing procedure (18). However, we use a more flexible semiparametric inference method, as described below in the next section.

### S1.3 Determine the number of latent factors *r*

To determine the number of unmeasured confounders *r*, one can use the joint-likelihood-based information criterion (JIC) (18). The JIC value is the sum of deviance and a penalty on model complexity:

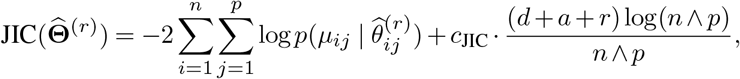

where 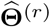 is the estimated natural parameter matrix with *r* unmeasured confounders and *d* + *a* observed covariates, and *c*_JIC_ *>* 0 is a universal constant set to be 1 by default.

Because “no unmeasured confounding” is not testable from observed data alone, we cannot certify that a dataset is unconfounded, but can provide actionable diagnostics that (a) flag when confounding is likely and (b) summarize how much it could matter for inference. This can be achieved by estimating the latent structure to quantify residual shared variation not explained by *X*, and summarize severity by the magnitude of change in gene-level effects when adjusting for 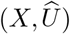 versus *X* alone (e.g., correlation or median absolute change of 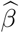 across genes). If the estimated number of latent factors is near zero and effect estimates are essentially unchanged after including 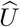, this provides evidence that residual unmeasured confounding beyond *X* is limited (while not constituting a proof of absence).

### S1.4 Comparison with reference-based confounder adjustment methods

Another approach to adjust for the unmeasured confounders is to utilize the information from negative control genes. This includes scMerge (54), RUV-III-NB (32) and RUVSeq (22) etc. These methods require users to specify a set of negative control genes, such as housekeeping genes, which are assumed to be solely due to unwanted variation between the two cells. The approach necessitates strong prior knowledge to accurately identify negative control genes, which may not always be available, especially in less well-characterized biological systems. This reliance on prior knowledge can limit the applicability of the method in novel or poorly understood contexts.

### S1.5 Comparison with RUV and RUV-III-NB

The primary reason for RUVr’s poor performance in this causal inference setting is its two-stage estimation strategy. RUVr first removes the variation associated with treatment *A* (and other observed covariates) and then searches for latent factors in the remaining residual variation. By design, this procedure identifies factors of unwanted variation that are orthogonal to treatment *A*. This is fundamentally at odds with the goal of adjusting for confounding in observational studies. A confounder *U* is, by definition, a variable that influences both the treatment *A* and the outcome *Y*. RUVr’s approach systematically fails to estimate any part of a latent factor that is correlated with the treatment, leading to underestimated variance and inflated Type I error.

This conceptual limitation is compounded by a model mismatch. RUVr assumes a linear model, which is ill-suited for the sparse, non-normal count data characteristic of single-cell sequencing even after log-transformation. In contrast, our method utilizes a more appropriate GLM framework and estimates effects iteratively, allowing it to identify confounders that are correlated with the treatment.

### S1.6 Relationship to data integration

Popular data integration methods aim to align datasets in a shared low-dimensional space to improve downstream tasks like clustering, visualization, and label transfer (19). These objectives differ from causal effect estimation: integration optimizes removal of variation that impedes cross-batch mixing, whereas causal DE requires adjusting for confounding that is correlated with treatment while preserving the treatment signal. When batch structure (or other unwanted variation) is aligned with treatment assignment, “removing batch” can also remove part of the treatment effect, so integration success metrics need not correspond to unbiased gene-level effects. In contrast, causarray treats technical and biological unwanted variation as potential confounding and adjusts for it through a generalized factor model coupled with doubly robust estimation, producing gene-wise causal estimands with valid uncertainty quantification under stated identification assumptions.

## Supplementary Information S2: Statistical inference

### S2.1 Counterfactual

#### Potential outcomes framework

Let 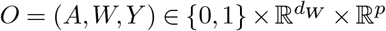 be a tuple of random vectors, where *A* is the binary treatment variable (e.g., presence or absence of a disease or perturbation), *W* is the vector of covariates (e.g., biological or technical factors influencing both treatment and outcome), and *Y* is the observed outcomes, defined as *Y* = *AY* (1) + (1 − *A*)*Y* (0), where *Y* (1) and *Y* (0) are the potential outcomes under treatment and control, respectively.

The potential outcomes framework assumes that for each individual or observation, there exist two potential outcomes: one if the individual receives the treatment (*Y* (1)) and one if they do not (*Y* (0)). However, only one of these outcomes can be observed for each individual, depending on whether they were treated (*A* = 1) or not (*A* = 0). This framework allows us to define causal effects in terms of these unobservable potential outcomes.

To estimate causal effects, we rely on the following key assumptions:

*Assumption 1* (Consistency) The observed response is consistent such that *Y* (*a*) = *Y* | *A* = *a*.

*Assumption 2* (Positivity) The propensity score *π*_*a*_(*W*) := ℙ(*A* = *a | W*) ∈ (*ϵ*, 1 − *ϵ*) for some *ϵ* ∈ (0, 1*/*2).

The positivity assumption is also known as the overlap condition or the common support condition. It requires that the two functions ℙ(*A* = 1 | *X* = *x*) and ℙ(*A* = 0 | *X* = *x*) share the same support in confounder values *X* = *x* (corresponding to the overlap condition or common support condition). Violations of this assumption are typically diagnosed by inspecting whether estimated propensity scores are extremely close to 0 or 1, which suggests regions of the covariate space where treatment assignment is nearly deterministic.

*Assumption 3* (No unmeasured confounders) *A* ⫫ *Y* (*a*) | *W*, for all *a* ∈ {0, 1}.

Under these assumptions (Assumptions 1–3), the observed outcome *Y* is conditionally independent of the treatment *A*, given the covariates *W*. This allows us to estimate the expected potential outcome for gene *j* under treatment (*a* = 1) or control (*a* = 0) as:

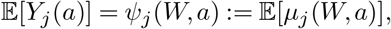

where *µ*_*j*_(*W, a*) = 𝔼 [*Y*_*j*_ | *W, A* = *a*] is a regression function that models the relationship between covariates, treatment, and outcomes.

Suppose we have a dataset 𝒟 = {*O*_1_, …, *O*_*n*_} consisting of i.i.d. samples from the same distribution as *O*. Let 𝔼_*n*_ denote the empirical measure over 𝒟, defined as:

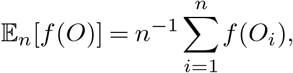

for any measurable function *f*. Equivalently, this computes the expectation with respect to the empirical distribution that puts weights 1*/n* on each observed data point. This represents the sample average of a function evaluated on all observations in the dataset.

A naive plug-in estimator for *ψ*_*j*_ can then be constructed by replacing the true regression function *µ*_*j*_(*W, a*) with its estimated counterpart 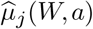 and using sample averages to approximate expectations. The resulting estimator is:

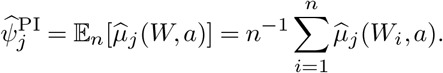

This plug-in estimator provides an estimate of the expected potential outcome by averaging predictions from the estimated regression model over all observations in the dataset.

While Assumptions 1–3 are foundational for causal inference, violations of the no unmeasured confounders assumption (Assumption 3) are common in real-world applications (18, 19). For instance, in single-cell transcriptomic studies, technical factors such as batch effects or biological heterogeneity (e.g., cell size or cell cycle stage) may act as unmeasured confounders. These unmeasured variables can bias estimates of causal effects by introducing spurious associations between treatment and outcome. Addressing this limitation motivates the need for methods that explicitly model and adjust for unmeasured confounders.

### S2.2 Target estimands

For semiparametric inference, a target estimand is a distributional functional of the observed random variables. For example, we can consider the average treatment effects (ATE), the standardized average treatment effect (SATE), the average treatment effect in levels or fold change (FC), and the LFC. Below, we define these estimands:

- 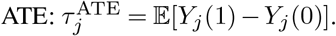
- 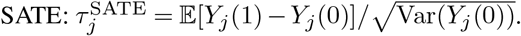
- 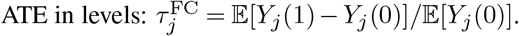
- 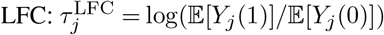

Here, we use *Y*_*j*_ to denote the random variable of the *j*th outcome and (*Y*_*j*_(0), *Y*_*j*_(1)) to denote its potential outcomes. Next, we present the corresponding influence functions under the identification assumptions, Assumptions 1–3. Before we present the influence functions, we introduce the uncentered influence function for 𝔼[*Y*_*j*_(*a*)] and 𝔼[*Y*_*j*_(0)^2^]:

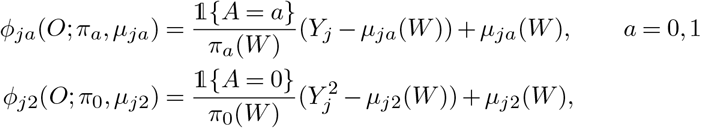

where *µ*_*ja*_(*W*) = 𝔼[*Y*_*j*_ | *W, A* = *a*] for *a* = 0, 1 and 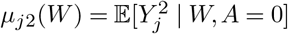. Note that the (centered) influence function of 𝔼[*Y* (*a*)] is given by *ϕ*_*ja*_(*O*; *π*_*a*_, *µ*_*ja*_) − 𝔼[*Y*_*j*_ (*a*)]. It follows that

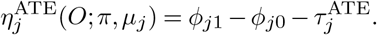

The efficient centered influence function of 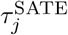 is given by

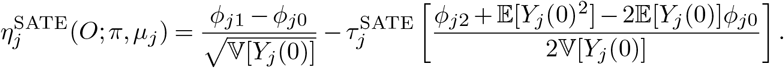

See for example, Equation (6) of (55) and Equation (4.3) of (8). Similarly, the efficient influence function of 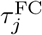 is given by

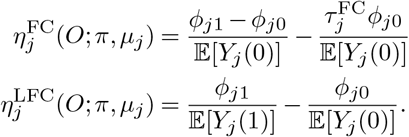

In the current paper, we restrict our focus to LFC; however, our implementation also allows the computation and inference using other estimands listed above. When computing the LFCs, we use the size-normalized counts *Y*_*ij*_ */s*_*i*_ adjusted by the size factors *s*_*i*_ in place of the raw count *Y*_*ij*_. This is akin to taking a weighted average of the sample to estimate ATE (and, subsequently, LFC). Otherwise, the effect will be driven by cells with large library sizes.

#### CATE

Under standard identification assumptions of consistency, conditional exchangeability, and positivity as in Assumptions 1–3, the conditional average treatment effect (CATE) is identified by *τ*_*j*_ (*w*) = *µ*_*j*1_(*w*) − *µ* _*j*0_(*w*). This also applies to conditional log-fold change. When one is only interested in the conditional effects in a subset of variables 𝒮 ⊂ [*d*_*W*_ + *a*], the DR-learner readily accommodates runtime confounding through the decomposition *τ*_𝒮_ (*w*) = 𝔼[*ϕ*(*O*) | *W*_𝒮_ = *w*_𝒮_]. This decomposition implies that one may estimate *τ*_𝒮_ (*w*) by regressing *ϕ*(*O*) on *W*_S_, i.e., modifying the final regression step of the DR-learner.

### S2.3 False discovery rate control

Genomic studies often involve testing thousands of hypotheses simultaneously, making it crucial to control statistical Type I errors. Two widely recognized error rate metrics are the Family-Wise Error Rate (FWER) and the False Discovery Rate (FDR), each suited to different contexts. Consider *p* hypothesis tests, let 𝒮 ⊂ {1,…, *p*} denote the set of discoveries, and ℋ_0_ ⊂ {1,…, *p*} denote the set of true null hypotheses. The false discovery proportion (FDP) is defined as the ratio of false positives to total discoveries:

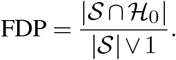

The FWER controls the probability of making at least one false discovery:

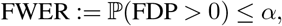

where *α* ∈ (0, 1) is a predefined significance level. This stringent control is particularly useful in scenarios where even a single false positive is unacceptable. However, FWER control often leads to reduced statistical power, especially in high-dimensional settings with many hypotheses, potentially overlooking true effects.

In contrast, FDR control provides a more balanced approach by controlling the expected proportion of false discoveries among all discoveries:

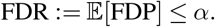

#### Algorithm S2

Multiple testing on standardized treatment effects

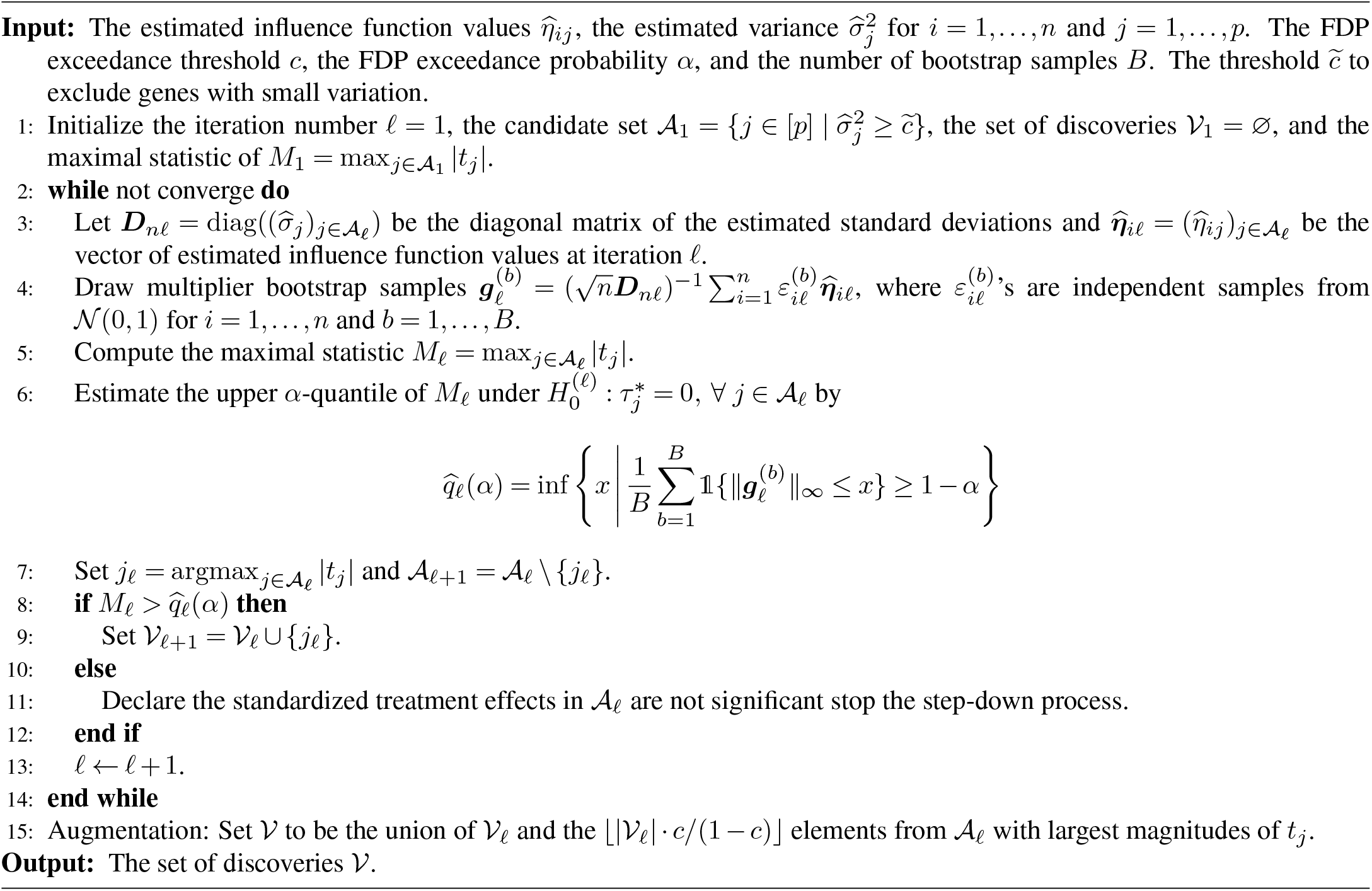

This approach enhances power in multiple testing scenarios and has become the standard for differential expression analysis in genomics due to its ability to identify more significant features while maintaining a low proportion of false positives (56). Importantly, FDR controls the *expected* proportion of false discoveries across repeated experiments but does not guarantee bounds on FDP in any single experiment. This distinction becomes critical in genomic studies where test statistics are often highly dependent, leading to variability in FDP across experiments.

To address limitations of standard FDR procedures, such as their inability to capture FDP variability in a single experiment, alternative error control metrics like False Discovery Exceedance (FDX) have been proposed:

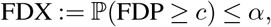

for a threshold *c* ∈ (0, 1). FDX provides stricter control by limiting the probability that FDP exceeds a predefined threshold *c*. This makes it particularly useful in applications where minimizing false positives is critical or when restricting analysis to a small subset of discoveries is desired.

To ensure robust error rate control tailored to genomic applications, causarray implements two complementary strategies for FDR control: (i) Benjamini–Hochberg (BH) Procedure: The BH procedure (56) is applied directly to P-values obtained from the semiparametric estimation framework. BH controls the FDR under independence or specific positive dependence structures among test statistics. (ii) Gaussian Multiplier Bootstrap: For tighter control of FDP variability, when test statistics are highly dependent, causarray incorporates a Gaussian multiplier bootstrap approach (Algorithm S2). It simulates null distributions to estimate FDP more accurately and provides robust FDR control even under complex dependence structures (8).

The choice between the BH and the Gaussian multiplier bootstrap depends on the dependency structure among test statistics. While BH is computationally efficient and widely used, it may not adequately control FDR under strong dependencies. The Gaussian multiplier bootstrap, on the other hand, accounts for complex dependency structures and provides more accurate bounds on FDP variability. Additionally, incorporating FDX offers an extra layer of conservatism for applications where minimizing false positives is critical. By offering these complementary strategies, causarray ensures robust error rate control tailored to diverse genomic applications while balancing power and error control.

## Supplementary Information S3: Data simulation and analysis

### S3.1 Data simulation

We consider two simulation settings. In the first simulation, we generate cells from zero-inflated Poisson distributions. In the second simulation, we use a specialized single-cell simulator Splatter (29) to generate cells with batch effects. Both simulations include 1 observed covariate and 4 unmeasured confounders. The details of the simulation are provided below.

#### Pseudo-bulk expression simulation details

The pseudo-bulk expression data are generated using a Poisson distribution with a zero-inflation component. The setup involves generating a latent signal matrix influenced by random noise and specific parameters. The data generation process is described in Algorithm S3 in detail. For experimental results in Fig. 2, we set *d* = 2 and *r*^*^ = 1, and vary *n* ∈ {100, 500, 1000, 5000}. For causarray, RUV, and RUV-III-NB, we provide the number of latent factors in *r* ∈ {2, 4, 6}. Because the simulated data consists of 3 cell types, which may be explained with 3 additional degrees of freedom, the best possible choice of the number of latent factors would be *r* = 4. The metrics for each experimental setting are calculated using 50 simulated datasets with varying random seeds.

##### Algorithm S3

Data generation process for pseudo-bulk gene expressions.

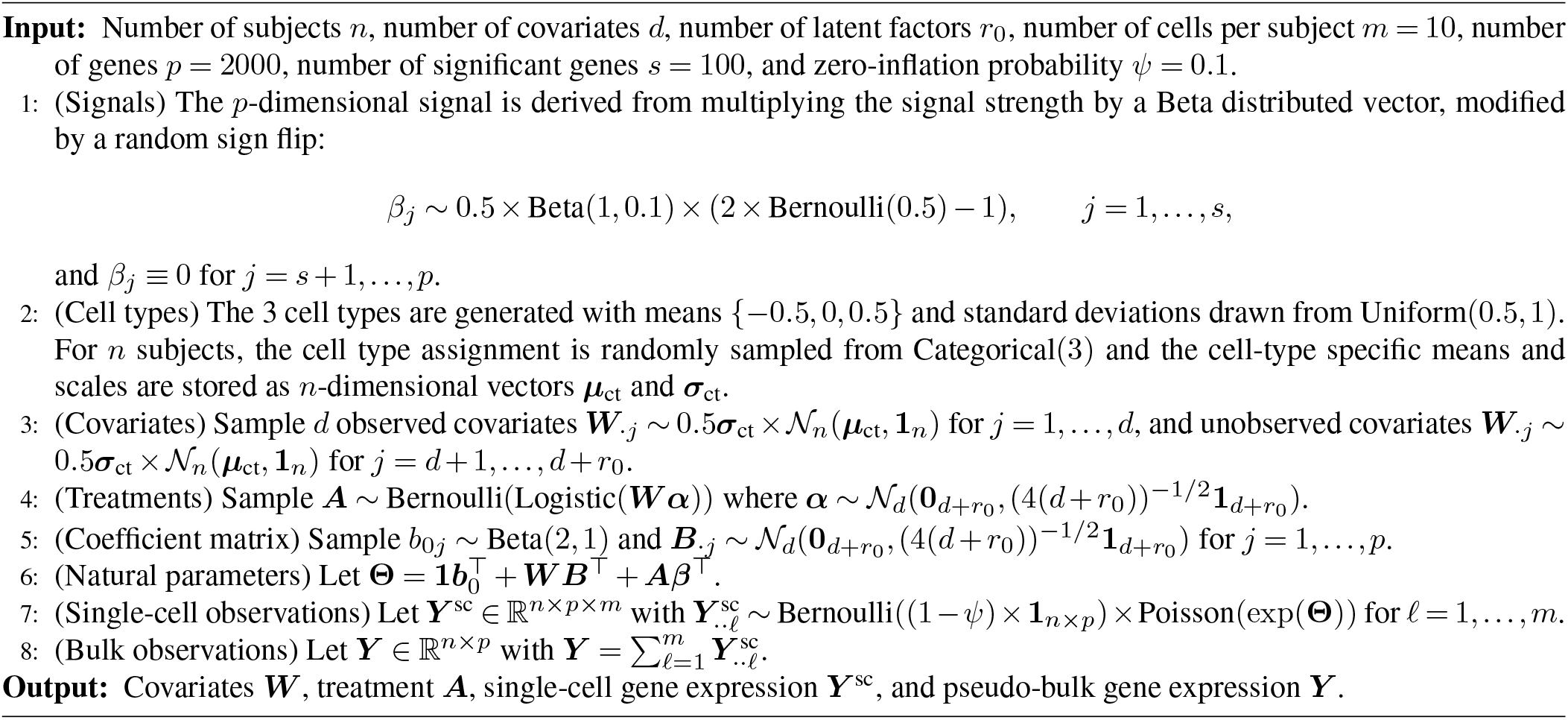

#### Single-cell expression simulation details

The single-cell expression data are generated by Splatter (29). Splatter explicitly models the hierarchical Gamma-Poisson processes that give rise to data observed in scRNA-seq experiments and can model the multiple-faceted variability. The data is generated from the `splatSimulate` function from the Splatter (1.26.0) package (29). When calling this function, the treatment effects are simulated with the parameters: `group.prob = c(0.5, 0.5), method = “groups”, de.prob=0.05`, `de.facLoc=1., de.facScale=0.5, de.downProb=0.5`; the dropout effects are simulated with the parameters: `dropout.type=“experiment”, dropout.mid=20, dropout` `.shape=0.001`; the batch effects are simulated with the parameters: `batch.facLoc=noise, batch.facScale=0.5`; while all the other parameters are the same as returned by the function `newSplatParams`. For experimental results in Fig. S2, we generate *d* = 1 covariates and *r* = 4 unmeasured confounders. We first generate (*d* + *r* + 1)*/*2 batches with equal sample sizes, which account for *d* + *r* degrees of freedom. To simulate varying confounding levels, we set `noise` in {0.1, 0.2, 0.3}. The metrics for each experimental setting are calculated using 50 simulated datasets with varying random seeds.

### S3.2 Benchmarking methods

To evaluate the performance of differential expression (DE) testing, we compare causarray with several established methods, both with and without confounder adjustment. Several linear methods, such as SVA (21), CATE (57), BConf (58), and dSVA (59), have been developed for confounder adjustment in bulk transcriptomic or DNA methylation data. Among these, CATE has demonstrated superior FDR control under linear models. However, such methods are less suitable for single-cell RNA-seq data due to sparsity and nonlinear mean–variance relationships. GCATE (18), based on a generalized linear model (GLM), outperformed CATE on both single-cell and pseudo-bulk data. As CATE, BConf, and dSVA perform similarly (60), we focus on methods better suited for single-cell data in this study.

Many methods aim to estimate confounding while performing single-cell treatment effect estimation, including Mixscape, contrastiveVI, cellOT, CPA, and PS (31, 50, 61–63). We compare causarray against CINEMA-OT and Mixscape as representatives of these approaches (optimal transport and matching).

Comparison methods included in simulations are grouped into two categories based on whether they account for unmeasured confounders.

Methods without confounder adjustment include:

- Wilcoxon rank-sum test: This nonparametric test is applied to deviance residuals obtained by regressing gene expression counts on measured covariates using a negative binomial GLM. The deviance residuals serve as input for the test, which does not explicitly account for unmeasured confounders.
- DESeq2 (30): This widely used method fits a negative binomial GLM to gene expression counts and adjusts for measured covariates. However, it does not account for unmeasured confounders, which may bias results in the presence of hidden variation.

Methods with confounder adjustment include:

- CoCoA-diff (R package `mmutilR 1.0.5`) (7): Designed for individual-level case-control studies, CoCoA-diff prioritizes disease genes by adjusting for confounders estimated from parametric models. We set `knn = 50` for cell matching, as suggested in the original publication, while keeping the remaining parameters unchanged. After adjusting for these confounders, the Wilcoxon rank-sum test is applied to the adjusted residuals, as recommended in the original paper.
- CINEMA-OT and CINEMA-OT-W (Python package `cinemaot 0.0.3`) (20): CINEMA-OT separates confounding sources of variation from perturbation effects using optimal transport matching to estimate counterfactual cell pairs. Compared to CINEMA-OT, CINEMA-OT-W further adjusts for differences in propensity scores between treated and control cells by matching before independent component analysis. We applied CINEMA-OT to library-size-normalized and log1p-transformed counts, setting the smoothness parameter to 1e-3 while maintaining all other parameters at their default values. Similar to CoCoA-diff, the Wilcoxon rank-sum test is applied to the adjusted residuals of CINEMA-OT or CINEMA-OT-W.
- Mixscape (31), a popular method based on matching for modeling perturbation effects. We applied the implementation from (20) to library-size-normalized and log1p-transformed counts.
- RUV-III-NB (R package `ruvIIInb 0.8.2.0`) (32): This method normalizes gene expression data using pseudo-replicates and a negative binomial model to remove unwanted variation induced by library size differences. The Kruskal-Wallis test (equivalent to the Wilcoxon test for two-group comparisons) is then applied to log-percentile adjusted counts, as suggested by the authors. However, RUV-III-NB does not directly adjust for library size and its ability to control FDR remains unclear, as it was not demonstrated in their experiments.
- RUV (R package `ruv 0.9.7.1`) (22): RUVr is used to estimate unmeasured confounders, which are then incorporated into DESeq2 for statistical inference based on both observed and estimated covariates. Before running RUV, we successively use the functions

`calcNormFactors`, `estimateGLMCommonDisp`, `estimateGLMTagwiseDisp`, and `glmFit` of edgeR package (4.0.16) (64) to extract residuals not explained by observed covariates and treatments.

This comprehensive benchmarking enables a thorough evaluation of each method’s ability to address unmeasured confounder estimation and perform robust statistical inference in simulated data settings. Users interested in applying these methods to new datasets may consider additional hyperparameter tuning to potentially improve performance. However, this typically requires defining a task-specific objective function and performing cross-validation. These steps are not straightforward in real-world applications where the true set of differentially expressed genes is unknown.

### S3.3 Evaluation metrics

To compare the performance of different methods, we use four evaluation metrics, focusing on two aspects: confounder estimation and biological signal preservation. DESeq2 and Wilcoxon are excluded from confounder estimation evaluation as they do not estimate unmeasured confounders or counterfactuals.

The performance of confounder estimation is assessed using two clustering-based metrics: Adjusted Rand Index (ARI) and Average Silhouette Width (ASW) (65). These metrics evaluate the quality of mixing in response and confounder spaces, respectively. Formally, measures the similarity between the clustering results based on the estimated control responses *Y* (0) and the true cell-type labels of the same samples. It adjusts for similarities that occur by chance:

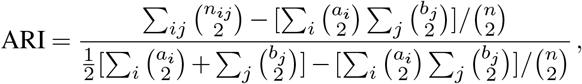

where *n* is the total number of samples, *n*_*ij*_ is the number of samples in both cluster *i* and partition *j, a*_*i*_ is the sum over rows in the contingency table, and *b*_*j*_ is the sum over columns. Higher ARI values indicate better conservation of cell identity based on estimated counterfactuals compared to true labels. When cell type is not the dominant confounder, as in our pseudo-bulk expression simulation, a larger ARI reflects better performance. In contrast, when cell type is the dominant confounder, as in the single-cell simulation, we seek to erase that signal, so a smaller ARI indicates more effective deconfounding. ARI ranges from −1 (complete disagreement) to 1 (perfect agreement), with 0 indicating random clustering. On the other hand, ASW quantifies how well each sample fits within its assigned cluster compared to other clusters. It is defined as:

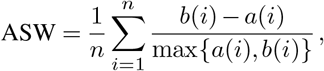

where *a*(*i*) is the average dissimilarity of sample *i* to all other samples within its cluster, and *b*(*i*) is the average dissimilarity to samples in the nearest neighboring cluster. ASW values range from −1 to 1, with higher values indicating better-defined clusters (65). For both metrics, median scores are scaled between 0 and 1 across methods within each simulation setup. For these two metrics, we use the implementations from the `scib` (`1.1.5`) package (65).

To evaluate biological signal preservation, we use False Positive Rate (FPR) and True Positive Rate (TPR), which are standard metrics derived from confusion matrices: PR quantifies the proportion of false positives among all true negatives:

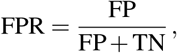

where FP and TN are false positives and true negatives, respectively. A lower FPR indicates fewer false discoveries relative to true negatives. Also known as sensitivity or recall, TPR measures the proportion of true positives among all actual positives:

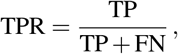

where TP and FN are true positives and false negatives, respectively. A higher TPR indicates better detection of true signals. These metrics provide complementary insights: FPR evaluates specificity by penalizing false discoveries, while TPR assesses sensitivity by rewarding correct detections. Together, they measure how well a method balances identifying true signals while avoiding false discoveries.

For the benchmarking methods presented in the paper, by design, the data are appropriate for either bulk or single-cell analysis, but not both. When many subjects have been measured, there is an intra-subject correlation among the cells. Analyzing at the pseudo bulk level circumvents this issue by treating subjects as the experimental unit. When the data is derived from a cell line or biological clones, the correlation between cells is negligible, and it is acceptable to use cells as the experimental unit. In this setting, a pseudo bulk analysis would not be advised. We describe the two simulation setups below.

### S3.4 Single-cell Perturb-Seq dataset

We utilize the Perturb-Seq dataset from (26), which enables high-resolution transcriptomic profiling of genetic perturbations in excitatory neurons. This scalable platform systematically investigates gene functions across diverse cell types and perturbation conditions, providing critical insights into neurodevelopmental processes (26). We focus on excitatory neurons of the dataset, a key population implicated in neurodevelopmental disorders such as autism spectrum disorders and neurodevelopmental delay, with perturbations targeting genes involved in neuronal development and synaptic function (26).

For preprocessing, if a given perturbation appears in fewer than 50 cells, all cells under that condition are excluded from the analysis. We then filter out genes expressed in fewer than 50 cells, resulting in a dataset containing 2926 cells under 30 perturbation conditions. The GFP (Green Fluorescent Protein) condition is used as a negative control to benchmark the effects of other perturbations by providing a baseline for comparison in downstream analyses. After filtering lowly expressed genes with a maximum count of fewer than 10, we retain 3221 genes.

The batch design is highly correlated with perturbation conditions; therefore, it is not included as a covariate in the model for testing. Instead, only the intercept is included as a covariate. For propensity score estimation, we incorporate the logarithm of library sizes as an additional covariate to account for technical variability and use GLM as the propensity score model. We performed a focused *Satb2* analysis to identify a biologically meaningful signal detected by causarray but missed by other correction methods, and then linked that signal to latent confounding. For *Satb2*, causarray identifies GO:0021953 (central nervous system neuron differentiation) with adjusted p-value 1.97e-05, while this term is absent in the RUV *Satb2* GO list. We then examined the genes underlying this GO term. The term contains 34 genes, including 26 genes classified as causarray-only vs RUV using the pre-specified criterion (causarray padj *<* 0.1, RUV padj *>*= 0.1 or missing), which represents biologically coherent genes preferentially identified by causarray (Fig. S5c). To characterize the confounder(s), we used the causarray latent factors 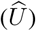 and computed their correlation with all genes, and the causarray-only set of DE genes shows substantial association with latent factors (Fig. S5d), consistent with the interpretation that these genes are influenced by unmeasured variation not explicitly modeled by standard alternatives. As one concrete example, *Hsp90ab1* in this GO term is strongly associated with latent confounding (maximum correlation = 0.419, best factor V7). Overall, these results support that causarray recovers biologically relevant Satb2 neurodevelopmental signals while explicitly accounting for latent confounding that can obscure or distort detection in other pipelines.

### S3.5 Single-nucleus Alzheimer’s disease dataset

This study integrates data from three single-nucleus RNA sequencing (snRNA-seq) datasets to investigate Alzheimer’s disease (AD): the ROSMAP-AD dataset (42) and two datasets from the Seattle Alzheimer’s Disease Brain Cell Atlas (SEA-AD) consortium (43), covering the middle temporal gyrus (MTG) and prefrontal cortex (PFC). These datasets provide complementary insights into AD pathology across different brain regions and donor cohorts.

The ROSMAP-AD dataset is derived from a single-nucleus transcriptomic atlas of the aged human prefrontal cortex, including 2.3 million cells from postmortem brain samples of 427 individuals with varying degrees of AD pathology and cognitive impairment (42). To ensure a balanced representation across subjects, we perform stratified down-sampling of 300 cells per subject, focusing on excitatory neurons while excluding two rare subtypes (‘Exc RELN CHD7’ and ‘Exc NRGN’). This preprocessing results in a dataset with 124997 cells and 33538 genes.

Next, we create pseudo-bulk gene expression profiles by aggregating gene expression counts across cells for each subject. Genes expressed in fewer than 10 subjects are filtered out, resulting in a final dataset of 427 samples and 26,106 genes. Binary treatment is defined based on the variable

‘age first ad dx’, which approximates the “age at the time of onset of Alzheimer’s dementia.” Covariates included in the analysis are ‘msex’ (biological sex), ‘pmi’ (postmortem interval), and ‘age death’ (age at death). Missing values for ‘pmi’ are imputed using the median of observed values.

The SEA-AD data are obtained from a multimodal cell atlas of AD developed by the Seattle Alzheimer’s Disease Brain Cell Atlas (SEA-AD) consortium (43). This resource includes snRNA-seq datasets from two brain regions: the middle temporal gyrus (MTG) and prefrontal cortex (PFC), covering 84 donors with varying AD pathologies.

For both MTG and PFC datasets, we perform stratified down-sampling of 300 cells per subject, focusing on excitatory neurons. Pseudo-bulk gene expression profiles are created by aggregating counts across cells for each subject. Genes expressed in fewer than 40 subjects are filtered out, resulting in final datasets with 80 samples and 24,621 genes for MTG and 80 samples and 25,361 genes for PFC. Covariates included in the analysis are ‘sex’, ‘pmi’, and ‘Age at death’. These variables account for biological and technical variability across donors.

To enable comparative analyses across the three datasets (ROSMAP-AD, SEA-AD MTG, and SEA-AD PFC), we restrict the analysis to 15586 common genes that are expressed in all three datasets. Genes with a maximum expression count below 10 among subjects are excluded to ensure robust comparisons.

#### Functional analysis

We further compare functional enrichment results between causarray and RUV using gene ontology (GO) terms associated with DE genes. Both methods identify overlapping top functional categories related to key biological processes associated with AD pathology (Fig. S6c). Both methods detect GO terms relevant to neuronal development and synaptic functions, which are critical for understanding AD pathology. However, causarray shows distinct enrichment in categories such as “positive regulation of cell development” and “negative regulation of cell cycle’, reflecting its increased sensitivity to synaptic and neurotransmission-related processes. In contrast, RUV’s results exhibit more dataset-specific enrichments, such as biosynthetic processes in SEA-AD (PFC), apoptotic processes in SEA-AD (MTG), and catabolic processes in ROSMAPAD (Fig. S6c). These findings suggest that causarray captures more generalizable biological signals across datasets. The visualization of the discovered networks, as defined as the top 5 GO terms and associated genes included in the top 100 DE gene discoveries, further highlights the enhanced sensitivity and comprehensiveness of causarray. Specifically, the causarray network contains 17 gene nodes and 81 edges, compared to 14 gene nodes and 57 edges in the RUV network (Fig. 4d). This greater interconnectedness in the larger causarray network suggests a more intricate and informative representation of underlying biological relationships, emphasizing its ability to capture broader and more relevant genetic factors associated with AD pathology.

## Supplementary Information S4: Extra results

### S4.1 Simulation

**Fig. S1.**
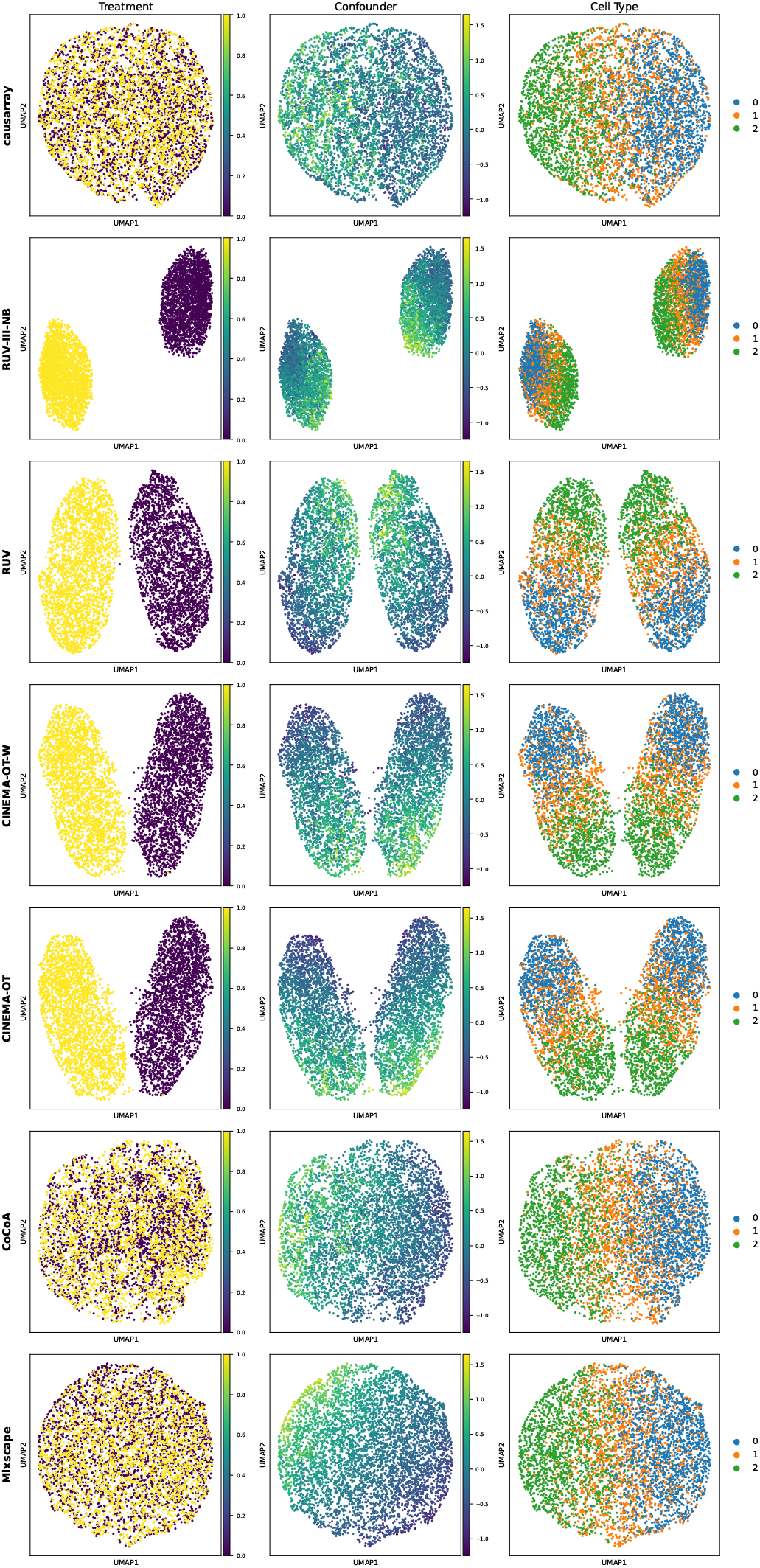
UMAP visualization of different methods in the confounder space on synthetic pseudo-bulk expression data with *n* = 5000 observations. The three columns are colored by treatment assignment (yellow for treated and purple for control), confounder value, and cell type. Because treatment is only probabilistically linked to the hidden confounder, treated and control cells should overlap in the actual confounder space. An estimator that recovers this geometry will therefore display intermingled colors in the UMAP, whereas clear color separation signals residual confounding or broken overlap. An ideal representation should eliminate treatment structure (mixing) while preserving both the continuous confounder gradient and the cell-type clusters. The visualisations show such a pattern for causarray, and CoCoA-diff. In contrast, other methods retain two clearly separated treatment clouds, probably due to the nonlinearity and sparsity nature of count data. This also explains the inflated false discovery rates of some of the other methods.

**Fig. S2.**
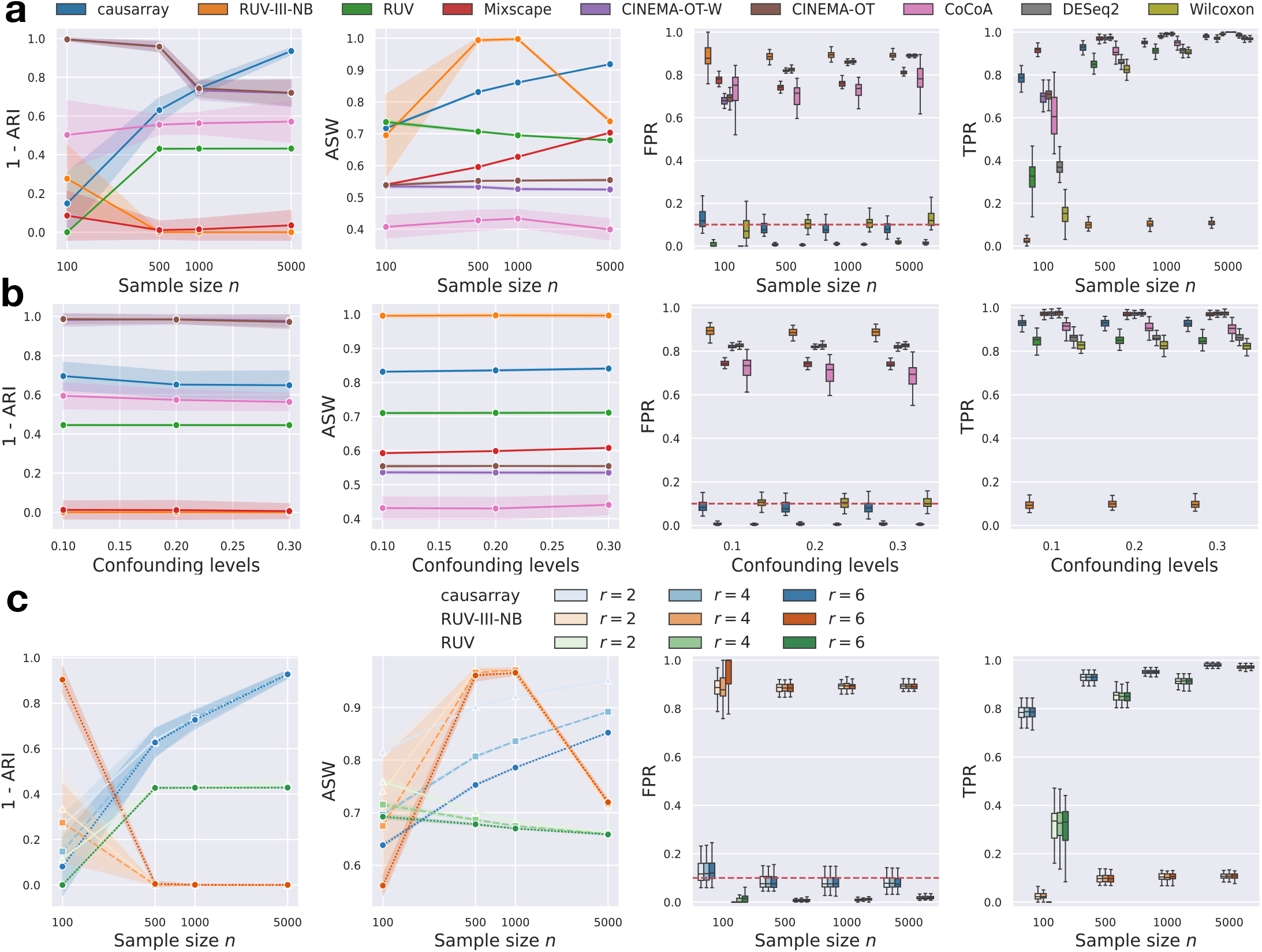
Benchmarking of causarray against other methods for single-cell differential expression testing on synthetic single-cell expression data under unmeasured confounders. The metrics for each experimental setting are calculated using 50 simulated datasets with varying random seeds. **a**, Line plots and box plots of different validation metrics for causarray and other methods with *r* = 4 latent factors and a moderate confounding level. Line plots show mean ARI and ASW scores for confounder estimation (shaded region represents values within one standard deviation). Because cell type is the dominant confounder, a smaller ARI indicates more effective deconfounding. Box plots (FPR, false positive rate, and TPR, true positive rate) indicate the performance of biological signal preservation. For box plots, the median is used as the center, the top and bottom hinges represent the top and bottom quartiles, and whiskers extend from the hinge to the largest or smallest value no further than 1.5 times the interquartile range from the hinge. **b**, Line plots and box plots of different validation metrics for causarray and other methods with varying confounding effects with *n* = 500 samples). **c**, Line plots and box plots of different validation metrics for RUV, RUV-III-NB, and causarray, with varying numbers of latent factors.

**Fig. S3.**
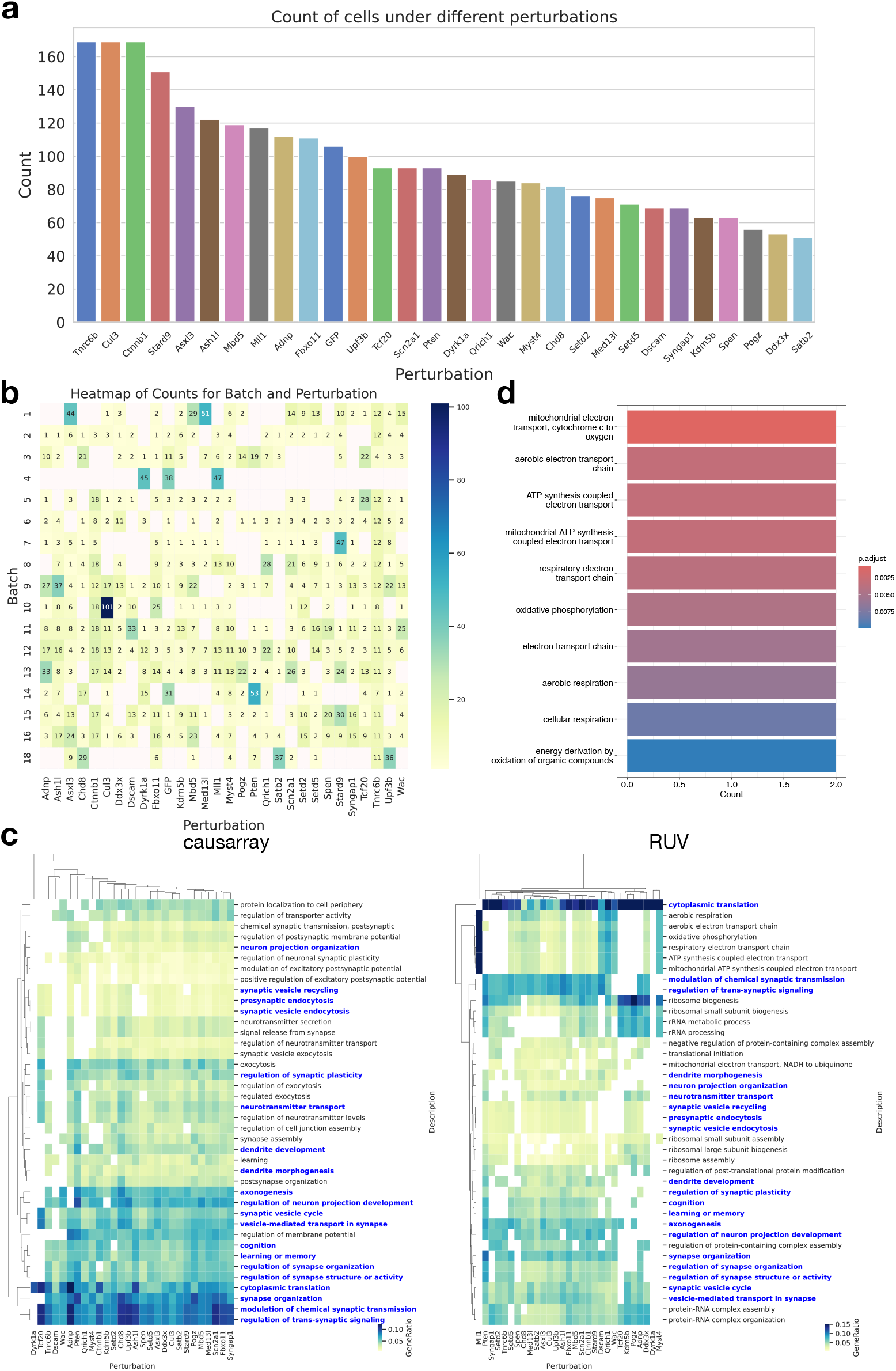
Additional results on the Perturb-seq dataset. **a**, Barplot of the number of cells in each perturbation. **b**, Heatmap of the number of cells in each batch and perturbation. The batch design and the perturbation assignment of the Perturb-seq dataset are highly correlated. **c**, Clustermaps of GO terms enriched in discoveries (FDR*<* 0.1) from causarray and RUV, respectively, where the common GO terms are highlighted in blue. Only the top 40 GO terms that have the most occurrences in all perturbations are displayed. **d**, Barplot of GO terms enriched in discoveries under *Mll1* perturbation from RUV.

**Fig. S4.**
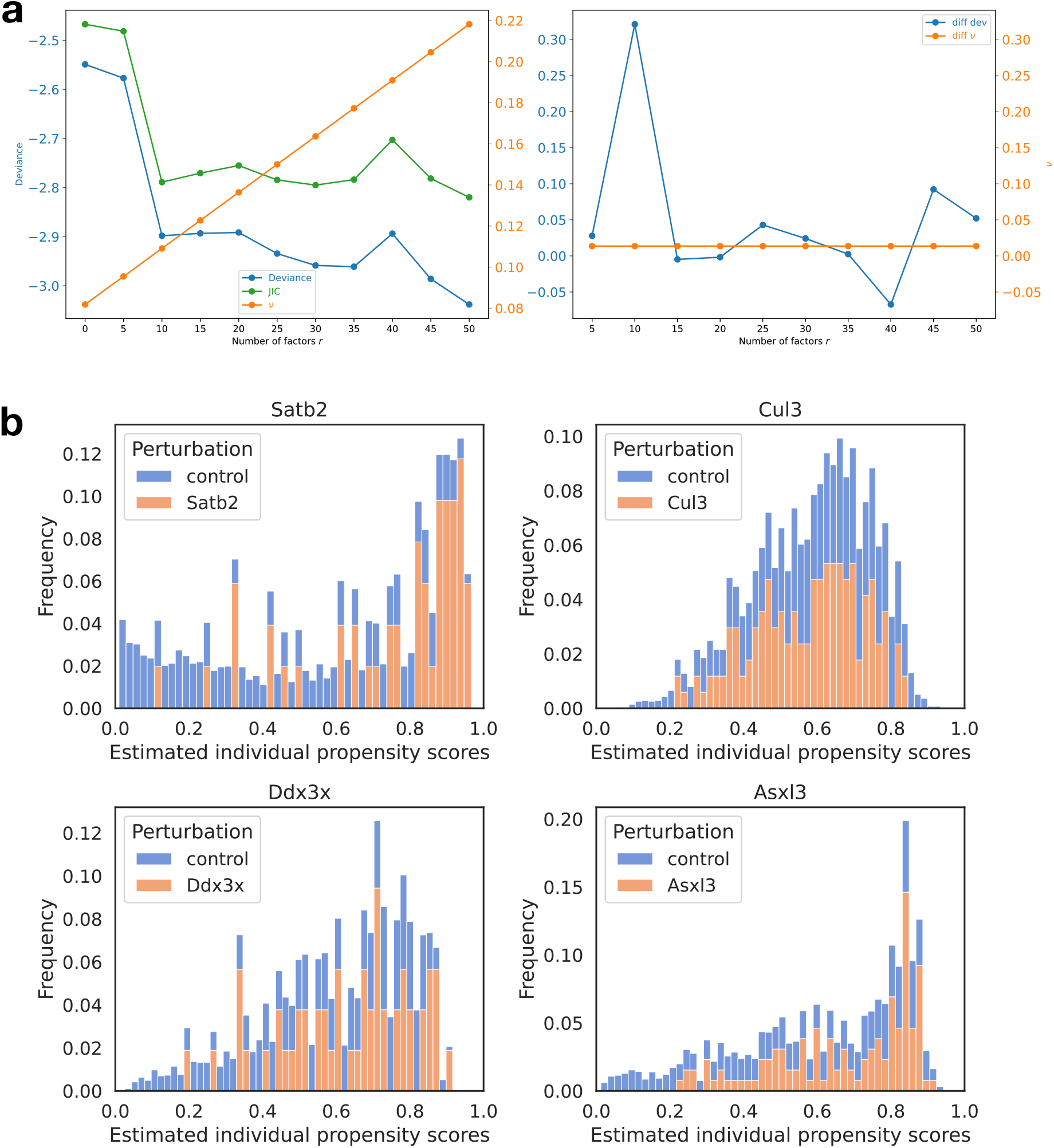
Estimation results of causarray on the Perturb-seq dataset. **a**, The JIC criteria suggest the number of latent factors *r* = 10. **b**, Histograms of estimated propensity score for the top 4 perturbations (*Satb*2, *Cul*3, *Ddx*3*x*, and *Asxl*3) with most significant genes (adjusted *P* value *<* 0.1).

**Fig. S5.**
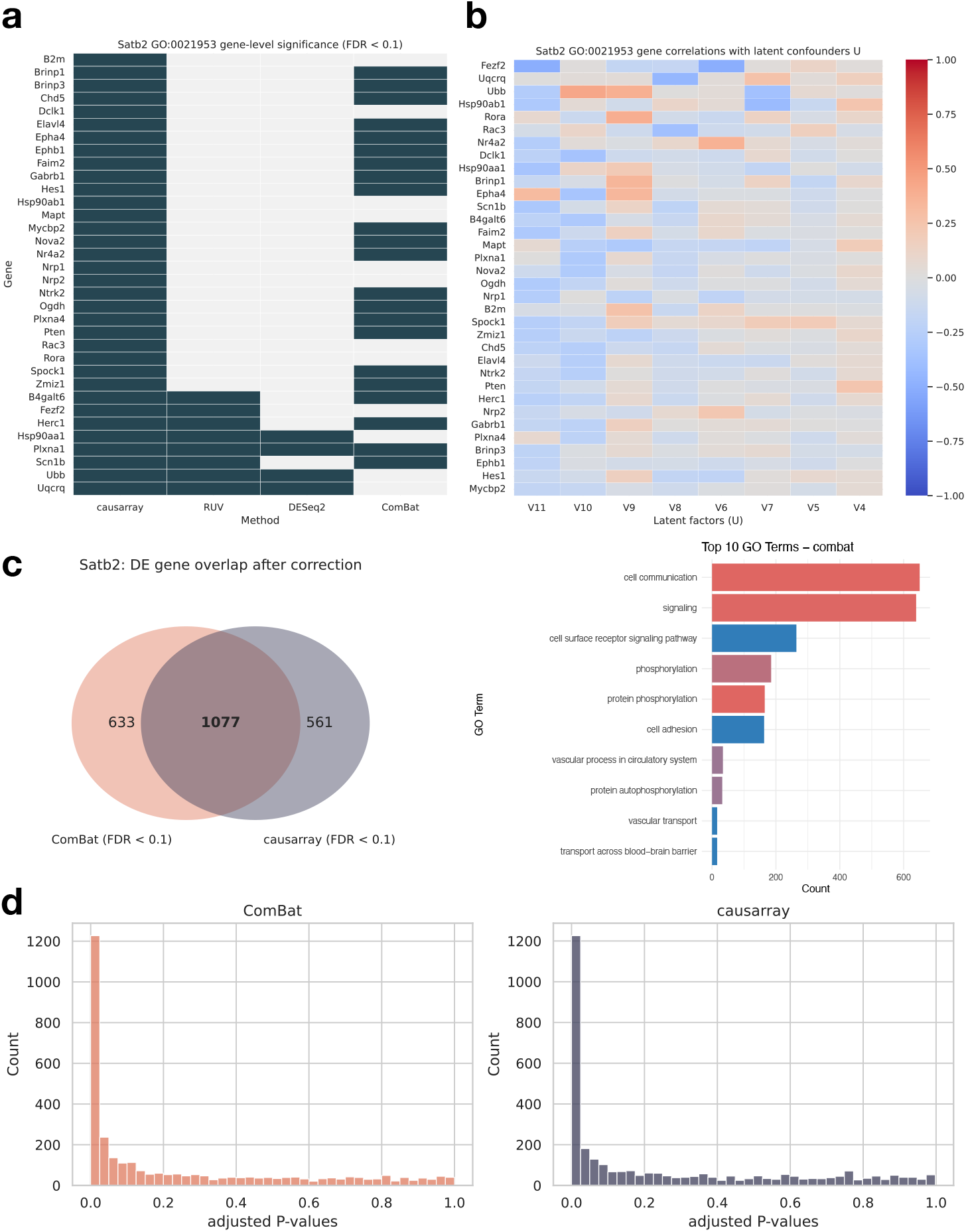
Extra results for *Satb2* perturbation on the Perturb-seq dataset. **a**, The DE genes by different methods that are associated with GO:0021953. **c**, The heatmap of correlation between the estimated latent factors by causarray and distinct DE genes by causarray versus RUV. **c**, The numbers of DE genes by ComBat and causarray, and the top 10 GO terms associated with ComBat’s DE genes. **d**, The distribution of adjusted P-values by ComBat and causarray.

**Fig. S6.**
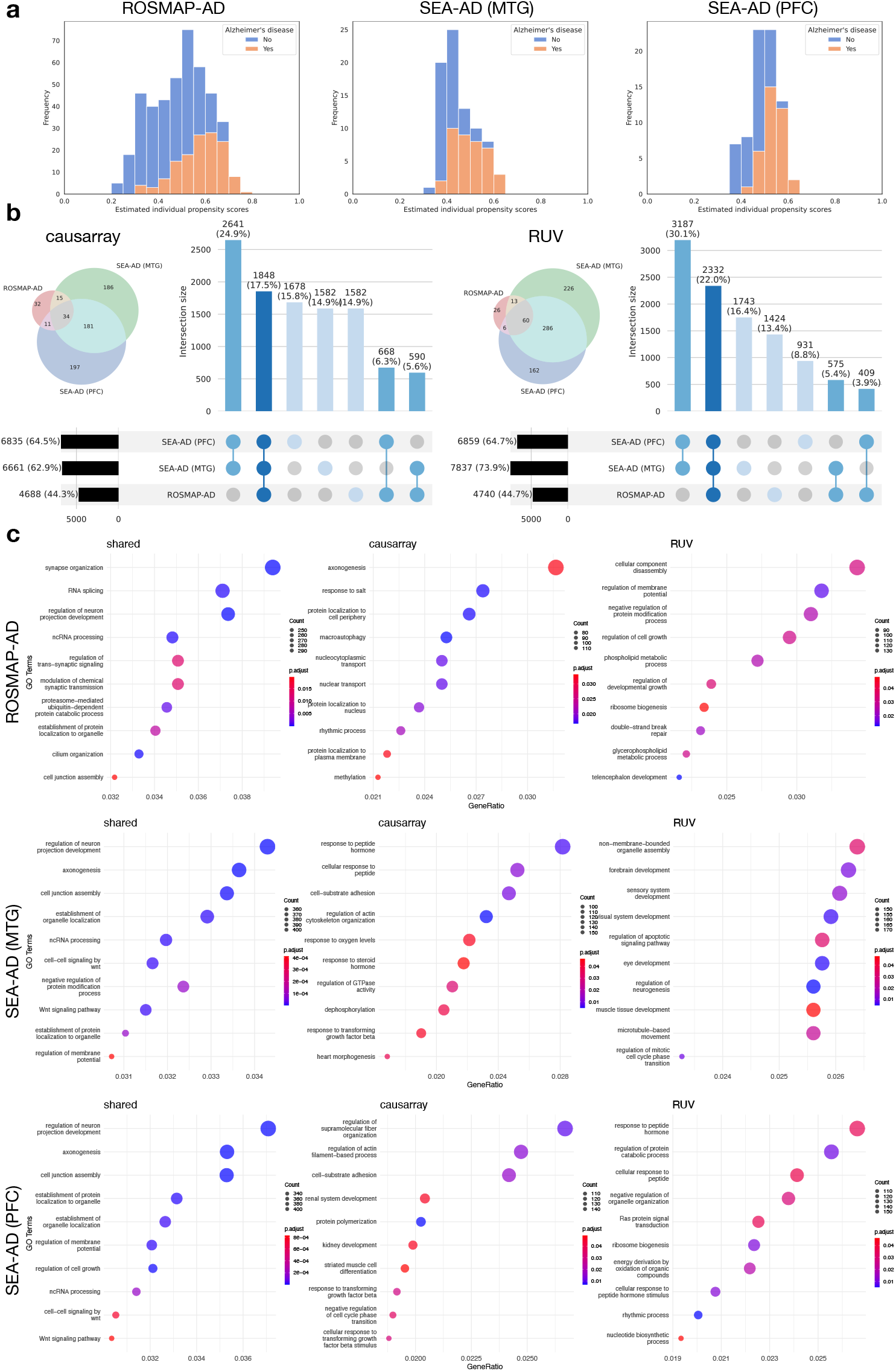
Extra experimental results in AD datasets. **a**, Histogram of estimated propensity score in three AD datasets. Note that the estimated scores remain within a moderate range away from 0 and 1, and the propensity score distributions overlap well, indicating no violation of Assumption 2. **b**, DE genes by causarray and RUV over 15586 genes (adjusted *P* value *<* 0.1). Venn diagrams show the associated GO terms (adjusted *P* value *<* 0.05, *q <* 0.2). **c**, Top gene ontology terms of the shared and distinct discoveries by causarray and RUV.

